# A biochemically-realisable relational model of the self-manufacturing cell

**DOI:** 10.1101/2021.06.07.447371

**Authors:** Jan-Hendrik S. Hofmeyr

## Abstract

As shown by Hofmeyr, the processes in the living cell can be divided into three classes of efficient causes that produce each other, so making the cell closed to efficient causation, the hallmark of an organism. They are the *enzyme catalysts* of covalent metabolic chemistry, the *intracellular milieu* that drives the supramolecular processes of chaperone-assisted folding and self-assembly of polypeptides and nucleic acids into functional catalysts and transporters, and the *membrane transporters* that maintain the intracellular milieu, in particular its electrolyte composition. Each class of efficient cause can be modelled as a relational diagram in the form of a mapping in graph-theoretic form, and a minimal model of a self-manufacturing system that is closed to efficient causation can be constructed from these three mappings using the formalism of relational biology. This Fabrication-Assembly or (*F,A*)-system serves as an alternative to Robert Rosen’s replicative Metabolism-Repair or (*M,R*)-system, which has been notoriously problematic to realise in terms of real biochemical processes. A key feature of the model is the explicit incorporation of formal cause, which arrests the infinite regress that plagues all relational models of the cell. The (*F,A*)-system is extended into a detailed relational model of the self-manufacturing cell that has a clear biochemical realisation. This (*F,A*) cell model allows the interpretation and visualisation of concepts such as the metabolism and repair components of Rosen’s (*M,R*)-system, John von Neumann’s universal constructor, Howard Pattee’s symbol-function split via the symbol-folding transformation, Marcello Barbieri’s genotype-ribotype-phenotype ontology, and Tibor Gánti’s chemoton.

## 1. Introduction

> “Organisms, cells, genes and proteins are complex structures whose relationships and properties are largely determined by their function in the whole. If an organism is our universe of discourse, *the cell is the star we gaze at* [emphasis added]: in its complexity and functionality even the simplest, tiniest cell dwarfs everything humankind has ever been able to engineer” (Wolkenhauer and Hofmeyr, 2007).

Being fragile, yet persistent; living longer than the functional lifetimes of one’s components: these hallmarks of life are made possible by the remarkable ability of living organisms to manufacture and individuate themselves autonomously as wholes, an ability that is arguably the most fundamental property that distinguishes living from nonliving systems. In order to grow, reproduce, metabolise, self-maintain, adapt and evolve, living organisms must first and foremost be able to self-manufacture. But, as the above quote (penned by Olaf Wolkenhauer) so poetically captures, if we want to understand organismal self-manufacture, we must surely first understand how the cell, the building block of all life as we know it, accomplishes this extraordinary feat.

Theories of life abound, so one would expect that by now, more than a century after the birth of biochemistry, we should be able to understand self-manufacture at the level of the modern cell’s molecular processes. However, the recent in-depth review by Cornish-Bowden and Cárdenas (2020) provides no evidence that any of the theories of life that directly address the question of cellular and organismal self-manufacture, of which Maturana and Varela’s (1980) theory of autopoiesis, Rosen’s (1991) theory of metabolism-repair or (*M,R*)-systems, and Gánti’s (2003c) chemoton theory are the most prominent examples, have found a satisfactory translation into cell biochemistry. Unfortunately the analysis by Hofmeyr (2017), which provides the basis for just such a translation, was for some reason not discussed in the review. The present article, using the context of (*M,R*)-systems and the formalism of relational biology as a starting point, builds on this analysis by developing a relational model of cellular self-manufacture that stands in a detailed modelling relation (Rosen, 1991) to cell biochemistry.

Why, after having used the term self-fabrication in previous publications (Hofmeyr, 2007, 2017), do I now rather use the term self-manufacture? *Manufacture* emphasises that at the molecular level the cell is a factory—as Barbieri (2005) so aptly pointed out, “life is artefact-making”. Manufacturing a composite artefact involves making from raw materials the parts of the artefact, which are then assembled into the artefact itself. Together these two separate processes, generally termed *fabrication* and *assembly* in the parlance of our manufacturing industry, manufacture the artefact from start to finish. So, whereas before I used fabrication as a synonym for manufacture, I need from now on to consign it to its rightful place as the first part of a manufacturing process. Other terms such as *production* (easily conflated with *reproduction*), *construction*, *making*, *maintaining* have also been used in this context, but they are not specific enough for my purposes.

The relational model developed in this article is a synthesis and further development of concepts that arose from Rosen’s (1991) metabolism-repair systems, from Louie’s further development in Louie (2009, 2013, 2017a) of Rashevsky (1954) and Rosen’s (1991) relational biology, and from Von Neumann’s (1966) universal constructor. The diagrammatic format of the model also allows the visualisation of key concepts that have helped build an understanding of what makes living organisms different from everything else: examples are Pattee’s (2001) epistemic cut and the symbol-function split via the symbol-folding transformation (Pattee, 1980) and Barbieri’s (1981) genotype-ribotype-phenotype ontology. I also use the model to highlight a serious deficiency in Gánti’s (2003c) chemoton theory.

Many theories of life aim to shed light on life’s origin, the ‘there’ in the question of ‘how did life get from there to here?”. What you are about to read is rather about the ‘here’, about my quest to reconcile the abstract concepts of Rosen’s relational biology with modern cell biochemistry.

## 2. Relational Biology

Relational biology studies the functional organisation of biological processes from a formal point of view. An excellent introduction to relational biology can be found in the Exordium of Louie (2009) and its complement in Louie (2017b). At its heart lies Rosen’s (1991) formalisation of Aristotle’s four so-called causes, the answers to the question ‘why something’ and therefore rather ‘becauses’, the parts that together make up a full explanation of that something. Their relation to each other is captured by a mathematical mapping *f* : *A* → *B*, more generally known in category theory as a morphism, which is an element in the hom-set *H*(*A, B*), the set of mappings from domain *A* to codomain *B*. The category **Set** of sets and mappings is the domain of relational biology: every process is a mapping such as *f* and every thing is a set such as *A* or *B* (Louie, 2009). The question ‘why *B*?’ is then answered by ‘because of the processor *f*, the *efficient* cause’ and ‘because of the *material* cause *A*?’ For the moment this suffices, but where *formal* and *final* causes fit in will become important further on. The mapping *f* : *A* → *B* can be depicted as a relational diagram in a graph-theoretic form (Fig. 1a) that was introduced by Rosen (1991) and refined by Louie (2009, 2013, 2017a). In Fig. 1a efficient causation is depicted by a dashed arrow and material causation by a solid arrow. In their graph-theoretic diagrams Rosen and Louie use solid-headed arrows and hollow-headed arrows, but I find the dashed and solid arrows easier to distinguish in more complicated diagrams.

**Figure 1:**
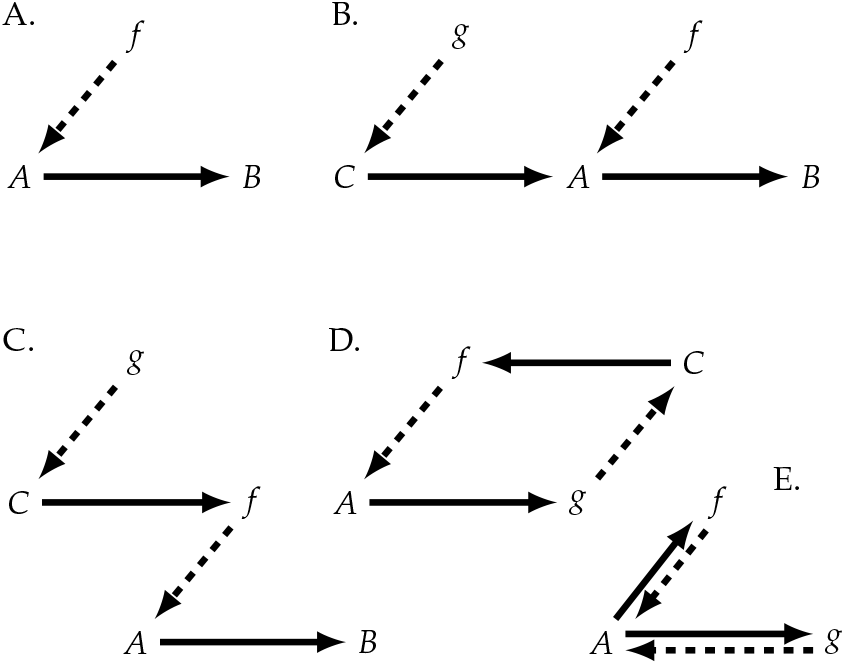
Graph-theoretic diagrams of mappings. (A) A single mapping. (B) Two mappings linked by a set *A* that is both the domain of *f* and the codomain of *g*. (C) A hierarchy of mappings in which *f* is functionally entailed by *g*. (D) A hierarchical cycle formed from the closure of the diagram in (C) by establishing a correspondence between *B* and *g*. (E) Equating *C* with *A* in diagram (D).

A key feature of category theory and, by implication, of relational biology is that mappings can be combined in various ways, such as where the codomain of one map is the domain of another (Fig. 1b); in the parlance of relational biology this forms a *path of material causation C* → *A* → *B*. However, nothing precludes the combination of mappings where a processor of one mapping is in the codomain of another mapping, as in Fig. 1c where *f* ∈ *H*(*A,B*), the codomain of *g*. Here we say that *f* is *functionally entailed* by *g* to form a *hierarchy of functional entailment*. The mappings in Fig. 1b can be composed into one mapping *f* ○ *g* : *C* → *B* (in category theory *f* ○ *g* means ‘*f* after *g*’), while the mappings in Fig. 1c compose into *g*(*c*) : *A* → *B* where *f* = *g*(*c*) with *c* ∈ *C*. In Rosen’s (1991) terminology Fig. 1a and b model *machines*, in which there is a clear distinction between ‘hardware’ *f* and *g* and ’software’ *A, B*, and *C*. Fig. 1c models a *mechanism*, in which this distinction is blurred: here *f* is both hardware and software. Machines form a proper subset of mechanisms. All three are models of what Rosen calls *simple* systems that can be computed (simulated) by a Turing machine.

**Figure 2:**
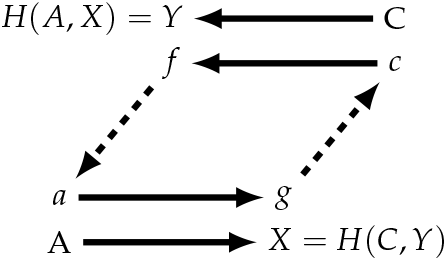
Impredicativity in an element-chasing version of the hierarchical cycle where a ∈ A, c ∈ C, f ∈ Y and g ∈ X. *X* = *H*(*C,Y*) and *Y* = *H*(*A,X*) so that *X* quantifies over the hom-set that contains *Y*, while *Y* quantifies over the hom-set that contains *X*. Substitution makes the impredicativity explicit: *X* = *H*(*C,H*(*A,X*)) and *Y* = *H*(*C,H*(*C,Y*)) so that *X* quantifies over a hom-set to which *X* itself belongs, and, similarly, *Y* quantifies over a hom-set to which *Y* itself belongs.

The cyclic diagrams in Figs. 1d and e form when the hierarchical diagram in Fig. 1c closes in on itself to form a *hierarchical cycle*; Louie (2009, Sec. 6.16) discusses the formal machinery that underlies such a closure. Unlike the other diagrams in Fig. 1 where one or more mappings are unentailed, both *f* and *g* in Figs. 1d and e are entailed within the cycle: *f* is the efficient cause of *g* and *g* is the efficient cause of *f*. Such diagrams (and by implication all systems modelled by them), where every efficient cause is entailed by another efficient cause in the diagram, are said to be *closed to efficient causation*, or, in short, *clef* (Louie and Poli, 2011). Figs. 1d and e model the simplest possible *clef* systems.^1^

In relational biology any system that realises a diagram that contains one or more hierarchical cycles is called a *complex system*; as long as some part of the system is *clef*, it is complex. If *every* efficient cause is entailed from within the complex system it is called an *organism*, thereby emphasising that being *clef* is a necessary condition for life; in fact, being *clef* is equivalent to autonomous self-manufacture, which, as noted in the introduction, is *the* fundamental property that distinguishes living from nonliving systems. Of course only a subset of all *clef* diagrams may qualify as models of real organisms as we know them; the aim of this article is to identify the most suitable one.

Hierarchical cycles are by their very nature *impredicative*. A definition of an object *X* is impredicative if it quantifies over a collection to which *X* itself belongs. Some impredicative objects, such as Russell’s famous ’set of all sets that are not members of themselves’ lead to logical paradoxes, while most others, such as ‘the smallest fish of all the fish in the pond’ are benign and not paradoxical. The situation in the hierarchical cycle is a bit more complicated than that in the definition above: here *X* quantifies over a set to which *Y* belongs, while *Y* quantifies over a set to which *X* belongs, making the cycle collectively impredicative (Fig. 2). Impredicativity implies non-computability (non-simulability) by a Turing machine in the sense that any algorithmic description of an impredicative system, when run on a Turing machine, either exists in a deadlock that cannot get going or in an endless loop that does not halt (Kercel, 2006; Louie, 2009). An everyday example would be me and my beloved wanting to go on a walk together. To get going she waits for me to start while I wait for her to start (a deadlock in which nothing happens). Now imagine that we somehow do get going, but I will only stop when she does and she will only stop when I do (we keep walking forever in an endless loop).

Having identified efficient cause with mappings *f* and *g* and material cause with sets *A* and *C*, we are now in position to fit *final cause* into the picture. Consider set *A* in Fig. 1b: its efficient cause is *g* and its material cause is *C*, respectively explaining *what produces A* and what *A is made from*. Final cause explains what *A is for*, its function in the diagram, which here is clearly to serve as material cause for *B*, i.e., in this diagram the final cause of *A* is *B*.^2^ Similarly, *B* is also the final cause of *f*—the function of *f* is to produce *B*. However, *A* is not only the material cause of *B*, it is also the final cause of *C* and *g*, a final cause that has been internalised in the diagram. In Fig. 1c *f* is not only the efficient cause of *B* but also the final cause of *g*, which again has been internalised in the diagram. In the *clef* diagram in Fig. 1d *g* is the final cause of *f* and *f* is the final cause of *g*; all final causes have been internalised in the diagram, making it autonomous not only because it is *clef* but also in the sense that it is its own final cause.

## 3. *Clef* metabolism-repair systems

One of Rosen’s central insights was that any model of a living cell must at the very least be able to account for the material transformations we call *metabolism*, as well as the *synthesis from the products of metabolism of the enzymes that catalyse the metabolic reactions*, which, as they lose their function with time and are degraded, have to be continually replaced from within the cell (Rosen, 1958a,b, 1959b, 1972, 1991). He called the latter process *repair* although the term ‘replacement’ is more appropriate in the context of the cell (Letelier et al., 2006). Fig. 3 shows how such a metabolism-repair or (*M,R*)-diagram can be constructed from a two-tier functional entailment hierarchy. Any relational model of the cell will have to contain such an (*M,R*)-substructure. Exactly which metabolic products *B* refers to is left to whoever is interpreting the diagram to decide. This issue will be resolved in Section 8.

**Figure 3:**
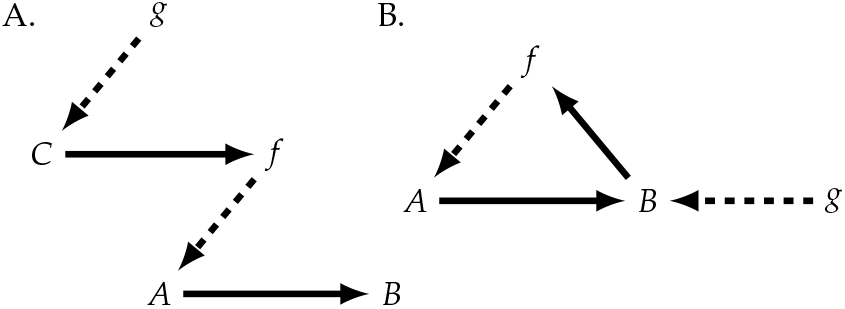
Forming a metabolism-repair system. (B) from the two-tier functional entailment hierarchy in (A) by identifying *C* with *B*. The implied biochemical interpretation of (B) is that the metabolic mapping *f* produces metabolic products *B* from nutrients *A*, while the repair mapping *g* produces enzyme catalysts *f* from *B*.

The diagram in Fig. 3b is, however, not closed to efficient causation: mapping *g* is still functionally unentailed. To be *clef* requires an additional mapping *h* that has a codomain of which *g* is an element and is itself identified with an existing entity in the (*M,R*)-diagram. This necessitates adding another level to the functional entailment hierarchy, as in Fig. 4a.

**Figure 4:**
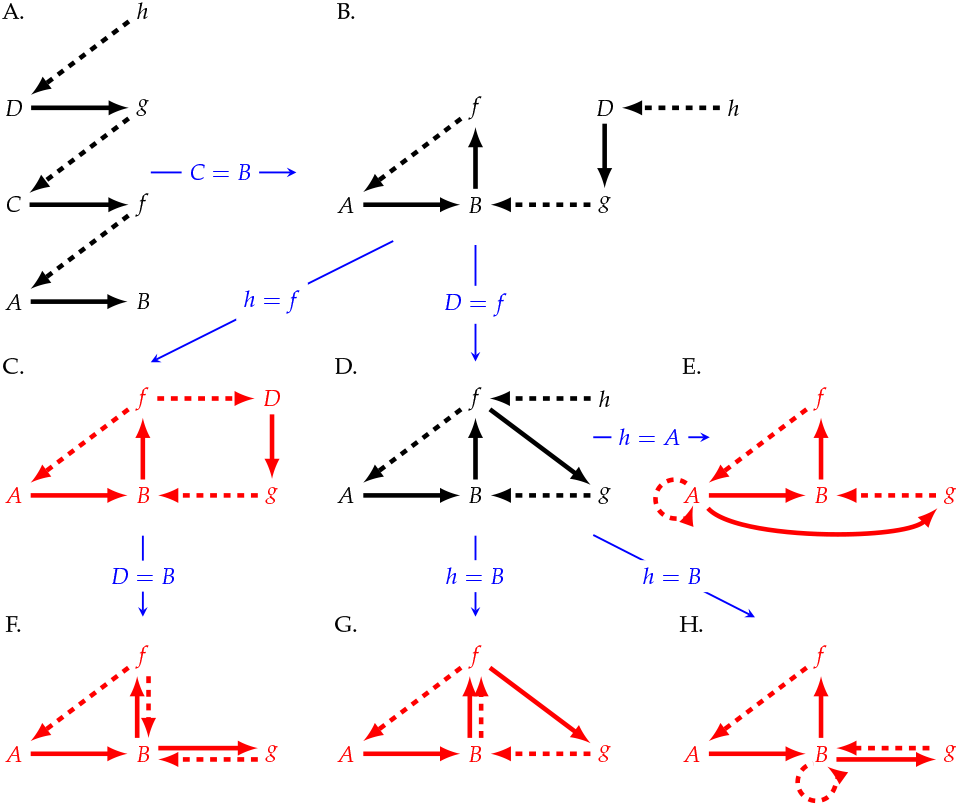
A. A three-tier functional entailment hierarchy. B-H. Stepwise identifications that lead to (F), the diagram that Rosen (1991, p. 238) constructed to exemplify a complex system, and (G), his replicative (*M,R*)-system (Rosen, 1991, p. 251). Diagrams (E) and (H) are alternative ways to (G) of embedding the ‘replication mapping’ *h* in (D) as discussed by Louie (2006, 2009). Although diagrams (C) and (E-H) are all closed to efficient causation, diagram (C), a diagram of mappings that models the closure of the three efficient causes in the living cell described by Hofmeyr (2017), is the central interest of this article.

The first step is then to create an (*M,R*)-substructure by identifying, as before, *C* with *B* (Fig. 4b). Rosen then chose to identify *D* with *f* to form Fig. 4d and *h* with *B* to form Fig. 4g, a system that he called a *replicative* (*M,R*)-system with the so-called ‘replication’ mapping *b* : *f* → *g* where *b* ∈ *B* (a rather unfortunate and misleading choice of name). More specifically, he found a way to place the identification of *h* with *b* on a solid mathematical footing.^3^

Despite the notorious difficulty of realising the replication mapping *h* as a cell process, this diagram has remained the target of all Rosen’s work on (*M,R*)-systems and that of his followers, critics and commentators. As such it has played a pivotal role in the development of relational biology and in anchoring category theory as the formal language of choice. However, apart from the fact that it has some congenial mathematical properties such as forming a Eulerian circuit (Louie, 2009), as a model of the cell nothing necessarily privileges it above other *clef* (*M,R*)-diagrams such as C, E, F, and H in Fig. 4. In fact, as far as models of the living cell and of organisms in general are concerned, I propose that the time has come to replace it with a model that actually stands in a full modelling relation to cellular processes in the sense that all mappings in the formal representation map onto (are realised by) real cell processes (Rosen, 1991).

In this article I demonstrate that, based on my analysis of the functional organisation of cell biochemistry (Hofmeyr, 2017), the diagram in Fig. 4c best fulfils the criteria for standing in a full modelling relation to the causal entailments in the living cell. It is ironic that its descendant Fig. 4f, which was used by Rosen (1991, p. 238) to exemplify a complex system, shares the properties that makes Fig. 4c a better model of the cell than Rosen’s replicative (*M,R*)-system in Fig. 4g. However, equating D with B turns out to be an undesirable constraint for the present analysis, so from here on I consider only Fig. 4c.

As Rosen noted for Fig. 4f, mapping *f* in Fig. 4c has two functions in the diagram, namely to make *b* = *f*(*a*) from *a* ∈ *A* and to make *g* = *f*(*d*) from *d* ∈ *D*. Were the diagram be assumed to model a mechanism, *f* should fractionate into the direct sum of *f*_1_ ∈ *H*(*A,B*) and *f*_2_ ∈ *H*(*D, H*(*B,H*(*A,B*))), which then has ineluctable implications either for *B* or for *g*. As will become clear in the next two sections, for my purposes I only need to consider the implication for *B*. Splitting *f* into the direct summands *f*_1_ + *f*_2_ requires *B* to split into the direct summands *B*_1_ + *B*_2_ (Fig. 5a). Mapping *g* can then be considered to be a monomorphism that maps *g* : *B*_1_ → *f*_1_ and *g* : *B*_2_ → *f*_2_ (Fig. 5a, diagrams 3 and 4). But this implies that *f*_1_ must split into *f*_11_ + *f*_12_ with *f*_11_ : *A* → *B*_1_ and *f*_12_ : *A* → *B*_2_ (Fig. 5a, diagrams 1 and 2), which in turn implies that *B*_1_ must split into *B*_11_ + *B*_12_ with *g* : *B*_11_ → *f*_11_ and *g* : *B*_12_ → *f*_12_, and so on. This leads to an infinite regress, which demonstrates that, were the diagram in Fig. 4c to model a natural system, that system cannot be a mechanism since it does not have a largest model.

**Figure 5:**
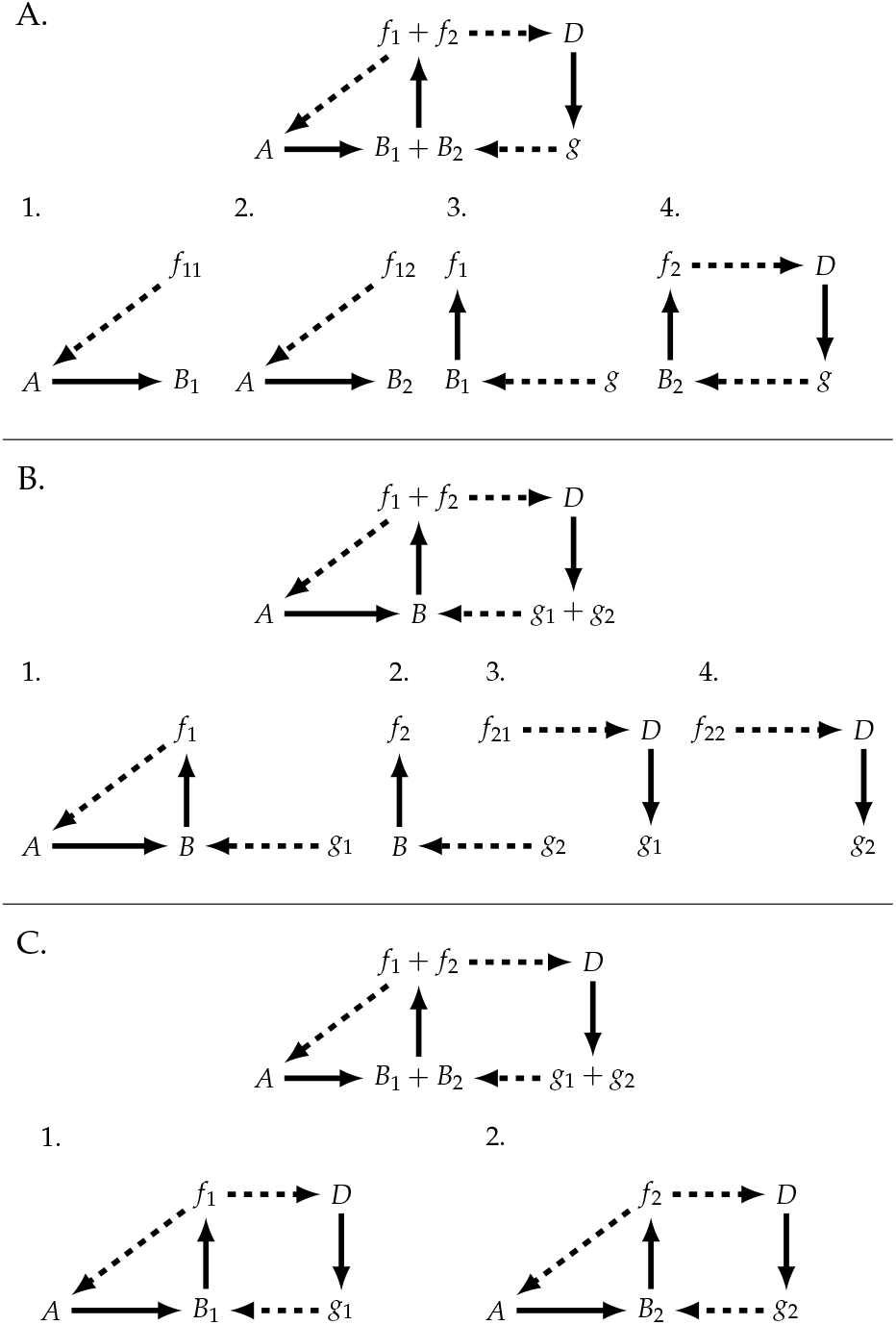
Consequences of assuming that the diagram in Fig. 4c models a mechanism. Mapping *f* in Fig. 4c splits into two direct summands *f*_1_ + *f*_2_, which requires either (A) set *B* to split into two direct summands *B*_1_ + *B*_2_, (B) mapping *g* to split into two direct summands *g*_1_ + *g*_2_, or (C) both *B* and *g* to split into respectively *B*_1_ + *B*_2_ and *g*_1_ + *g*_2_. The diagrams in (A), (B) and (C) are the direct sum of their subdiagrams. The text explains why in (A) *f*_1_ has to split into *f*_11_ + *f*_12_ and in (B) *f*_2_ has to split into *f*_21_ + *f*_22_.

This notwithstanding, I shall show that, by associating the *formal causes* of *B*_1_ and *B*_2_ with mapping *f*_1_, the regress can be stopped at the level shown in Fig. 5a, resulting in a *clef* diagram that is an amalgam of a mechanism (the sum of diagrams 1, 2 and 3) and a hierarchical cycle isomorphic to Fig. 2.

If *B* does not split into *B*_1_ and *B*_2_, the implication for *g* of splitting *f* into *f*_1_ + *f*_2_ is that *g* would have to split into the direct summands *g*_1_ + *g*_2_ with *g*_1_ : *B* → *f*_1_ and *g*_2_ : *B* → *f*_2_ (Fig. 5b, diagrams 1 and 2). In turn this forces *f*_2_ to split into *f*_21_ + *f*_22_ with *f*_21_ : *D* → *g*_1_ and *f*_22_ : *D* → *g*_2_ (Fig. 5b, diagrams 3 and 4), which in turn implies that *g*_2_ must split into *g*_21_ + *g*_22_ with *g*_21_ : *B* → *f*_21_ and *g*_22_ : *B* → *f*_22_, and so on. Again this leads to an infinite regress. In his corresponding treatment of Fig. 4f, Rosen (1991) claimed to have only considered the implication for *g*, but to arrive at diagrams 1 and 2 in Fig. 5c, the equivalents of the two diagrams in Rosen (1991, Fig. 9f.3), the implications for *B* and *g* would both have to be taken into account. In Fig. 5c the original diagram is the direct sum of diagrams 1 and 2. Repeating the argument for diagrams 1 and 2 leads to smaller and smaller models of Fig. 4c, clearly demonstrating that we are dealing with a Rosennean complex system and not a Rosennean mechanism.

## 4. Formal cause in graph-theoretic diagrams

Although formal cause is sporadically mentioned in relational biology, it has been conspicuous in its near absence from all discussions of (*M,R*)-systems to date. Despite, as we shall see, its pivotal role in self-manufacturing systems, there is to the best of my knowledge no example of its incorporation into (*M,R*)-diagrams. In fact, formal cause is discounted by some researchers as “less fundamental than the efficient cause and the material cause for understanding the nature of metabolic closure” (Cárdenas et al., 2018). A possible reason for this is proposed in the discussion in Section 12.

Formal cause, the fourth Aristotelean ‘because’, is the answer to the question ‘what is it *to be* something’ in the sense of being the *actualisation* of some *prior model* (the formal cause) of that something, model being used here in the broadest sense: a prior existing representation such as a mental or concrete image, an idea, a propensity, design, blueprint, template, program, mould, pattern or even a prior exemplar of that something (Alvarez, 2009; Falcon, 2019). For example, *to be* the sculpture of David is to be the actualisation in marble (material cause) of the representation (formal cause) in Michelangelo’s (efficient cause) head and his sketches of David.

Hofmeyr (2018) provided the first extensive treatment of formal cause in the context of the graph-theoretic diagrams of relational biology, and used it show that Rosen’s (1959a) purported paradox in Von Neumann’s (1966) description of kinematic self-reproducing automata is avoided by the appropriate incorporation of formal cause. As explained next, formal cause is associated with either efficient cause or with material cause.

In the first case the processor of a mapping is a combination of efficient and formal cause, in the sense that the formal cause ‘informs’ or ‘programs’ the efficient cause (analogously one can also consider, as did Kercel (2006), the formal cause to be a constraint on the efficient cause). One configuration (Fig. 6a) is that the formal cause *σ* parameterises the efficient cause *f* to form an *informed* or *programmed* efficient cause *f_σ_*, which is then a single entity. A biological example of Fig. 6a would be an enzyme that catalyses (efficient cause) the reaction in an active site on the enzyme, the chemical configuration of which determines which particular substrate (material cause) the enzyme binds and which reaction it catalyses. The active site can therefore be interpreted as not only the site of catalysis, but also a prior model (formal cause) that is actualised as the product of the reaction.^4^ In another configuration (Fig. 6b) the efficient and formal causes are distinct entities *f* and *σ* that combine into a processor complex; being distinct allows them to be domains, co-domains, or processors of other mappings. For want of a better word I shall call these formal causes *freestanding*. A biological example of Fig. 6b would be the combination of an mRNA molecule (freestanding formal cause) with a ribosome (efficient cause) which together produce a polypeptide (see Hofmeyr (2018) for an extensive discussion of these and of non-biological examples).

**Figure 6:**
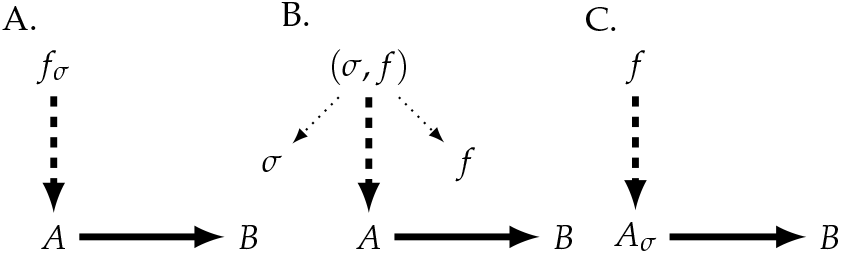
Formal cause in graph-theoretic diagrams. (A) Formal cause *σ* of *B* is associated with efficient cause *f* by parameterisation to *f_σ_* (a single entity with intrinsic formal cause). (B) Formal cause *σ* of *B* combines with efficient cause *f* to form the pair (*σ, f*), which is an element of the Cartesian product Σ× {*f*}, where Σ = {*σ*_1_, *σ*_2_, …}. The dotted arrows are projection maps that allow *f* and *σ* to appear as distinct entities in the diagram. (C) Formal cause *σ* of *B*, the propensity of *A* to transform into *B*, is intrinsic to material cause *A*.

Fig. 6c shows the second case, in which the formal cause is an intrinsic property of the material cause.^5^ Instead of informing the efficient cause, formal cause now informs the material cause. In an uncatalysed, reversible chemical reaction A ⇌ B the formal cause of B would be the intrinsic propensity of A to transform into B, while that of A would be the intrinsic propensity of B to transform into A; in a sense the formal cause of B can be thought of as a ‘model’ of B inherent in A, and *vice versa*. Biological examples would be the folding of a one-dimensional string of amino acids (a polypeptide) into a three-dimensional protein, or the self-assembly of different protein subunits into a multimeric protein complex, such as the two *α* and two *β* subunits into haemoglobin. In these examples the information for how to fold and to self-assemble is provided by the aminoacid sequence itself and by the chemical properties of the protein subunits: the formal cause is intrinsic to the material cause. However, correct folding and self-assembly can only take place in a specific chemical context, here a watery environment with buffered pH and strictly controlled ionic strength and electrolyte composition in which the molecules move around and collide through Brownian motion. Hofmeyr (2017) called this context the *intracellular milieu*, but these conditions can of course also be recreated in a test tube. Changing the context can cause misfolding and mis-assembly or even prevent these processes. For example, adding 6M urea or replacing water with chloroform unfolds (denatures) and disassembles protein complexes. Here the context clearly performs a crucial function, and can be equated with the efficient cause of these processes.^6^ One could object that the context does not play an active enough role to be considered an efficient cause, but this is incorrect. The major driving force for folding and self-assembly of polypeptides is the entropic force of the hydrophobic effect, which is the tendency of non-polar molecules or parts of molecules to aggregate in a watery environment, not by attracting each other but by being pushed together by water molecules. The hydrophobic effect minimises the area of contact between water and non-polar molecules, in so doing maximising the entropy of the water. Furthermore, the pH, ionic strength and electrolyte composition determine the state of dissociation and solvation of the functional groups on proteins and nucleic acids, properties that play an important part in the specificity of self-assembly (McManus et al., 2016; Hofmeyr, 2007, 2017). This third incarnation of formal cause will come into play in the model of a self-manufacturing system developed in the next section.

Fig. 6b and its description above glossed over an important facet of such a configuration, namely that the efficient cause must be able to translate, interpret or decode the information embedded in the formal cause into a form that it can execute. Consider the situation where you, a language X speaker, wants to assemble a composite object bought in kit form but the instructions are in language Y; or an aspiring Morse code operator that has lost the paper on which the translation from dots and dashes to letters of the alphabet are written; or a living cell with defective aminoacyl-tRNA synthetases that have lost the ability to couple the correct amino acids to their corresponding anticodon-carrying tRNAs. In all these cases a code is missing, where a code is defined as a set of arbitrary rules (the code mapping) selected from a potentially unlimited number in order to establish a specific correspondence between two independent worlds (Barbieri, 2015). Fig. 7 shows how the decoding of formal cause and the code mapping can be incorporated in the graph-theoretic diagrams of relational biology. This is discussed extensively in Hofmeyr (2018).

**Figure 7:**
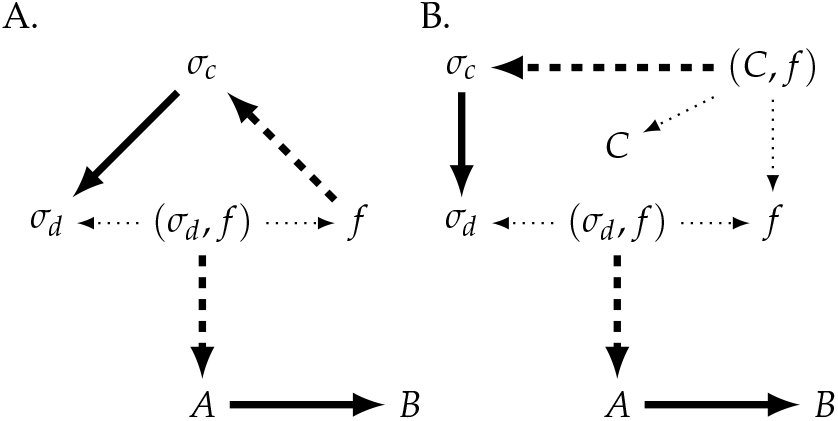
Decoding the formal cause. (A) Efficient cause *f* decodes an encoded formal cause *σ_c_* into *σ_d_*, which forms the processor (*σ_d_, f*) in association with *f*. Here the code mapping is implied. (B) The code mapping object C combines with *f* to form the decoding mapping (*C,f*), which then decodes *σ_c_* into *σ_d_* (Hofmeyr, 2018).

## 5. Manufacture = Fabrication + Assembly

As a point of departure for developing an understanding of the mechanistic aspect of Fig. 5a (diagrams 1, 2, and 3), Fig. 8 describes an abstract model of a manufacturing system in which a fabricator produces from raw materials a set of components from which a composite object is assembled by an assembler. This model borrows the essentials of Von Neumann’s (1966) concept of a universal kinematic constructor, but breaks it down into separate fabrication and assembly processes. As noted in the introduction, real-life manufacturing of composite objects usually proceeds in this way.

**Figure 8:**
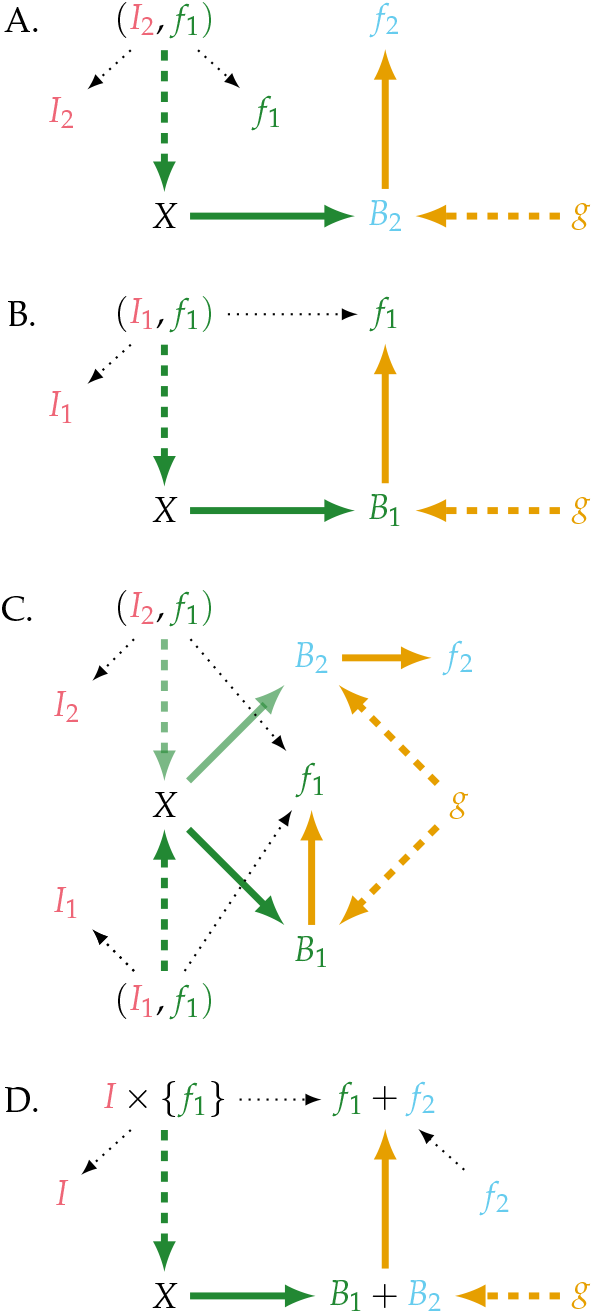
A generic fabricator-assembler manufacturing system. (A) Given a set of descriptions *I*_2_, fabricator *f*_1_ makes from materials *X* a set of components *B*_2_ which are then assembled by *g* into a composite automaton *f*_2_. (B) Here *f*_1_ fabricates its own component set *B*_1_ from *X* given its own description set *I*_1_. Assembler *g* then assembles *f*_1_ from *B*_1_. (C) The combination of (A) and (B). (D) A simplified form of diagram (C) that will be used in the rest of the article; *I* is the set of description sets, here {*I*_1_, *I*_2_}, the formal causes of *B*_1_ and *B*_2_. A dotted arrow is either a projection from a Cartesian product or from an element of a Cartesian product, an injection into a direct sum, or a composition of a projection and an injection, e.g., from *I* × {*f*_1_} to *f*_1_ + *f*_2_ via *f*_1_.

Consider as a modern-day example of such a system a 3D printer (the fabricator) that, when provided with a set of descriptions produced by slicer software, prints the set of plastic components of some composite object from a raw material such as the commonly-used polylactic acid fibre (PLA). In a separate process the components are assembled into the object by an assembler (Fig. 8a). In principle, one could also envisage the 3D printer printing its own component set that is then assembled into a daughter copy of the printer (Fig. 8b). One may be tempted to claim that the 3D printer has replicated itself, as indeed is done for the groundbreaking and award-winning 3D printer RepRap, which is described on its wiki page^7^ as “humanity’s first general-purpose self-replicating manufacturing machine”, followed by:

> “RepRap takes the form of a free desktop 3D printer capable of printing plastic objects. Since many parts of RepRap are made from plastic and RepRap prints those parts, RepRap self-replicates by making a kit of itself—a kit that anyone can assemble given time and materials. It also means that—if you’ve got a RepRap—you can print lots of useful stuff, and you can print another RepRap for a friend.…”

On closer inspection the claim that RepRap has replicated itself is at best disingenuous, and can only be based on a severely impoverished version of the concept of self-replication. All that RepRap can do in reality is print a subset of its components, but even if it were possible for it to print all of its components, it still cannot assemble itself—for that an external agent is needed (“… a kit that *anyone* can assemble…”). This is the situation in Fig. 8b, where, given the set of descriptions *I*_1_, fabricator *f*_1_ can produce only its own set of components *B*_1_, which then needs to be assembled into a new copy of *f*_1_ by assembler *g*. Were the original *f*_1_ to become dysfunctional, this new copy of *f*_1_ would replace it.

There is, however, something unexplained in Fig. 8. How does assembler *g* ‘know’ how to assemble both *f*_1_ from *B*_1_ and *f*_2_ from *B*_2_? Just as *f*_1_ needs descriptions to fabricate a component set, so *g* needs instructions on how to assemble a set of components into its corresponding object. It seems that one must therefore either assume a *universal assembler g* that needs to be able to follow the assembly instructions (associate with formal causes) for any number of different objects, or one must assume *dedicated assemblers*, which would here be *g*_1_ and *g*_2_, each having been trained (informed by a specific formal cause) to assemble one particular object by internalising its assembly instructions.

Let us reflect for a moment on the meaning of ‘universal’ in what I have just called a universal assembler. Von Neumann (1966) used this adjective to describe an automaton, the universal constructor, that can construct every other automaton, including itself. The abstract fabricator *f*_1_ described in Fig. 8 is universal in that, given the appropriate set of descriptions, it can produce any set of components, even its own. The abstract assembler *g* is universal in that, provided with the appropriate assembly instructions, it can in principle assemble any object from its component set, even itself. If universal assembler *g* can assemble both *f*_1_ and itself then the system would be self-manufacturing.

There is, however, no necessity for invoking the notion of an ‘informed’ assembler, albeit universal or dedicated. The manufacturing system as described in Fig. 8 with an ‘uninformed, untrained’ assembler *g* can be salvaged by invoking the formal cause configuration in Fig. 6c, in which the information for self-assembly of the object is intrinsic to the members of its component set. The assembler *g* is then the enabling context that drives the self-assembly of *f_i_* from the components in *B_i_*. Such a context would be difficult, but not impossible, to imagine for the macro-scale at which RepRap operates, but is eminently feasible at the nanoscale where molecules move, collide and react freely through Brownian motion. Gánti (2003a) used the term *fluid automaton* and *fluid machinery* to describe this situation.

The mappings in Fig. 8a and 8b are identical to diagrams 1 and 2 in Fig. 5a, with *X*, (*I*_1_, *f*_1_) and (*I*_2_, *f*_1_) equal to *A, f*_11_ and *f*_12_. The fabricator-assembly system in Fig. 8d can now be closed to efficient causation by assigning to *f*_2_ the function of mapping a material cause *D* into the assembler mapping *g*. The resulting diagram in Fig. 9a can be recast into Fig. 9b, the non-regressive form of Fig. 5a that incorporates formal cause *I*. It is now a simple matter to extend *X* → *B* to *A* → *X* → *B* by adding a new formal cause *I*_3_ that associates with *f*_1_ to form the mapping (*I*_3_, *f*_1_) : *X* → *B*_3_; *B*_3_ is then converted by *g* into mapping *f*_3_ : *A* → *X* (Fig. 9c). Composition of mappings *f*_3_ and *I* × {*f*_1_} yields Fig. 9d. Finally, with *B* = (*B*_3_ + *B*_1_) + *B*_2_ and *f* = (*I* × {*f*_1_} ○ *f*_3_)+*f*_2_, we have recaptured our starting diagram Fig. 4c.

**Figure 9:**
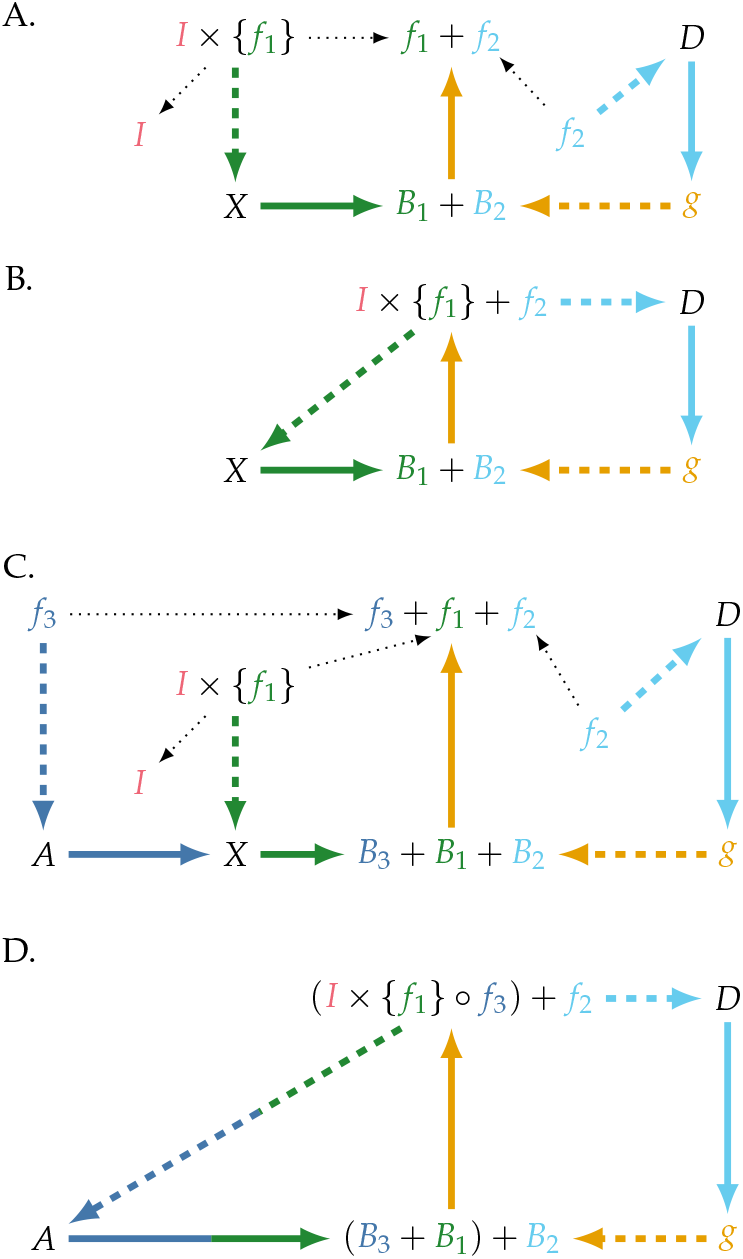
(A) Closure to efficient causation of Fig. 8d by designating *f*_2_ as the efficient cause of *g* through the mapping *f*_2_ : *D* → *g*. (B) is a compact form of diagram (A), and can be seen to be the non-regressive form of Fig. 5a through the explicit incorporation of the freestanding formal causes *I* = {*I*_1_, *I*_2_}. (C) shows how providing *f*_1_ with an additional description set *I*_3_ so that *I* = {*I*_1_, *I*_2_, *I*_3_} augments diagram (A) with an additional mapping *f*_3_ : *A*→*X*. (D) Composing *f*_3_ and (*I* × {*f*_1_}) into a single mapping *I* × {*f*_1_}○*f*_3_ yields a compact diagram that retains the relational structure of Figs. 5a and 4c.

The main difference between Fig. 9a and 9b (and between Fig. 9c and 9d) is the degree of visibility of formal cause *I*. In Fig. 9a and 9c *I* is explicit as a freestanding formal cause, whereas in Fig. 9b and 9d it is embedded in the mappings *I* × {*f*_1_} + *f*_2_ and (*I* × {*f*_1_}○*f*_3_) + *f*_2_. This obscuring of information and the resultant invisibility of information in the simple *clef* diagrams of Fig. 4 may be one of the main reasons why so little progress has been made in their biological realisation. What Figs. 8 and 9 make clear is that a fabricator-assembly manufacturing system must have a persistent source of freestanding information that is not destroyed by its use; for this to be an integral part of a *clef* system, the information must be materially embodied and internalised. As discussed in section 4 this inevitably leads to a requirement for a decoding and translation subsystem. This will become clear in the relational cell model that builds on the diagram in Fig. 9c.

Note that all three types of formal cause are present in Fig. 9: the subscripts 2 and 3 in *f*_2_ and *f_3_* represent *σ* in Fig. 6a, *I* represents the freestanding *σ* in Fig. 6b, and the subscripts 1, 2 and 3 in *B*_1_, *B*_2_, and *B*_3_ can be considered referents to *σ* in Fig. 6c.

Although I have used examples from biology, art and technology to illustrate and illuminate, nothing in this and the previous section depends on these examples. What I have formally shown is how to arrest the regress caused by splitting *f* into two direct summands *f*_1_ and *f*_2_ and the subsequent requirement for *B* to split into the direct summands *B*_1_ and *B*_2_ at the level of Fig. 5a. This was made possible by incorporating the formal causes for fabrication products into Fig. 5a by associating them with a fabricator mapping *f*_1_, and by embedding the formal causes for self-assembly products into the elements of sets *B*_1_ and *B_2_*. Doing so has of course not turned the diagram in Fig. 5a from a Rosennean complex system into a Rosennean mechanism; the existence of the hierarchical cycle precludes that. Being *clef*, Fig. 5a and, by implication, Fig. 4c are relational models of an organism, which clearly is neither a mechanism nor a machine.

## 6. Modelling cellular self-manufacture

The diagram in Fig. 9a serves as the core upon which to build the biochemically-realisable model of the selfmanufacturing cell, and will from here on be called the Fabrication-Assembly system or, in short, the (*F,A*)-system. In principle all that needs to be done is to provide additional formal causes to extend Fig. 9a with more mappings, such as in Fig. 9c, until we arrive at a relational model (the diagram in Fig. 11) that is congruent with Fig. 10, which is a view of the biochemical pathways that underlie cellular self-manufacturing. Hofmeyr (2017) provided all the biochemical detail that led to this view and is best read in conjunction with the present article. The construction of the extended (*F,A*) cell model in Fig. 11 has been the main aim of this article. However, is just as important to relate the (*F,A*) cell model back to Fig. 4c, our starting point in the land of simple *clef* systems. This is done in Fig. 12, which shows the three sets of efficient causes that correspond to the three mappings in Fig. 4c, repeated here as the left-hand diagram inserted above the cell model. The top right diagram is the (*F,A*)-system in Fig. 9c, which was the starting point for the development of the full (*F,A*) cell model. These diagrams clearly show that both the (*F,A*)-system and (*F,A*) cell model are open to material causes (*A* and *D*; nutrients and electrolytes) and open to the freestanding formal causes (*I* = [*I*_1_, *I*_2_, *I*_3_}; DNA and mRNA), while being closed to efficient causes (*f*_1_, *f*_2_, *f*_3_ and *g*; enzymes, transporters, and ribosomes).

**Figure 10:**
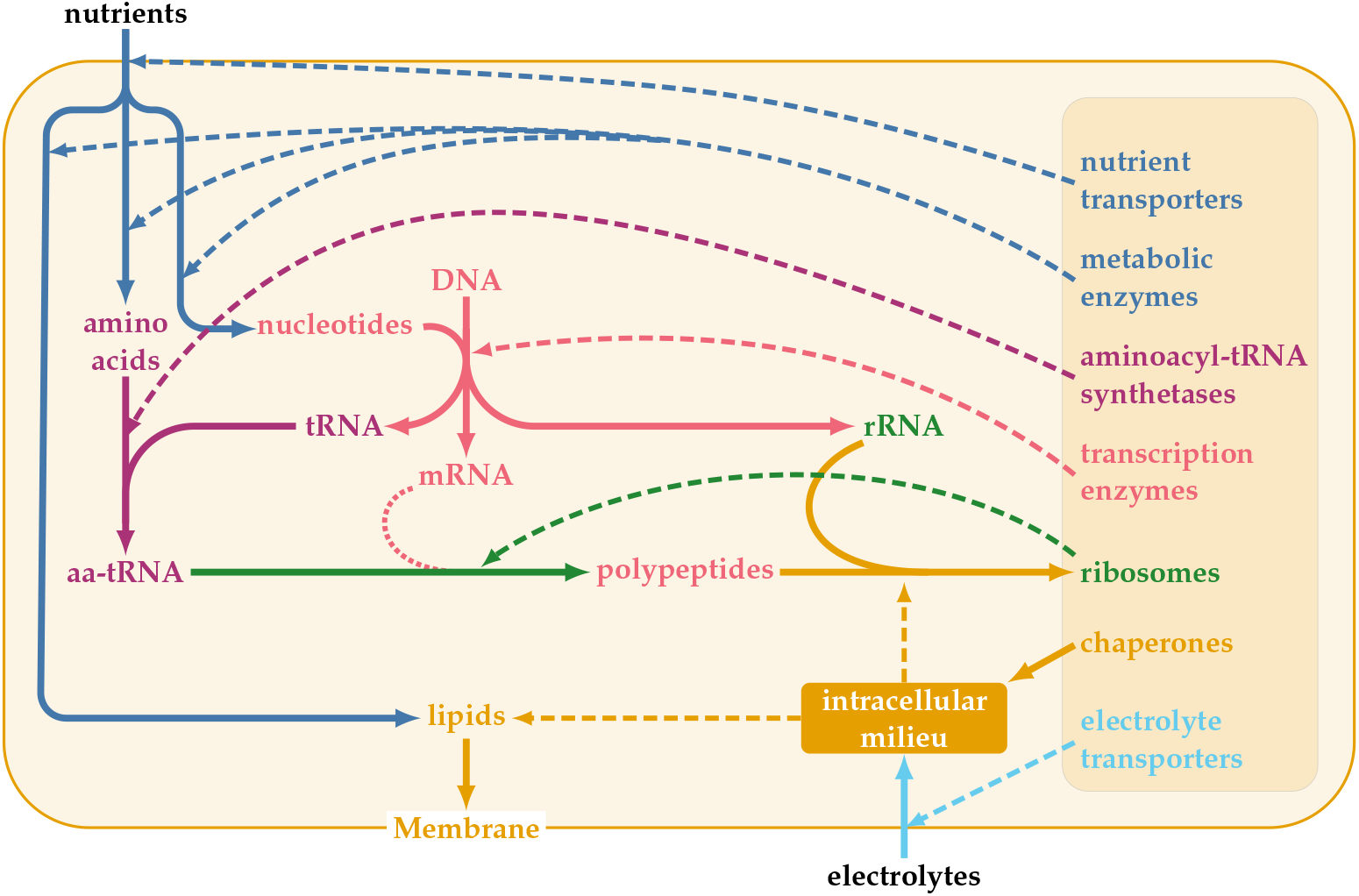
The functional organization of the biochemical processes that underlie the self-manufacture of the cell (adapted from Hofmeyr (2017)). The dashed arrows emanate from the efficient causes, while the solid arrows emanate from material causes. The dotted line emanating from mRNA depicts formal causation by transfer of sequence information from DNA to the amino acid sequences in polypeptides. Note that, similar to DNA and mRNA, the yet-to-be folded polypeptides are coloured red to emphasise that they are sequence information-carrying material symbols.

**Figure 11:**
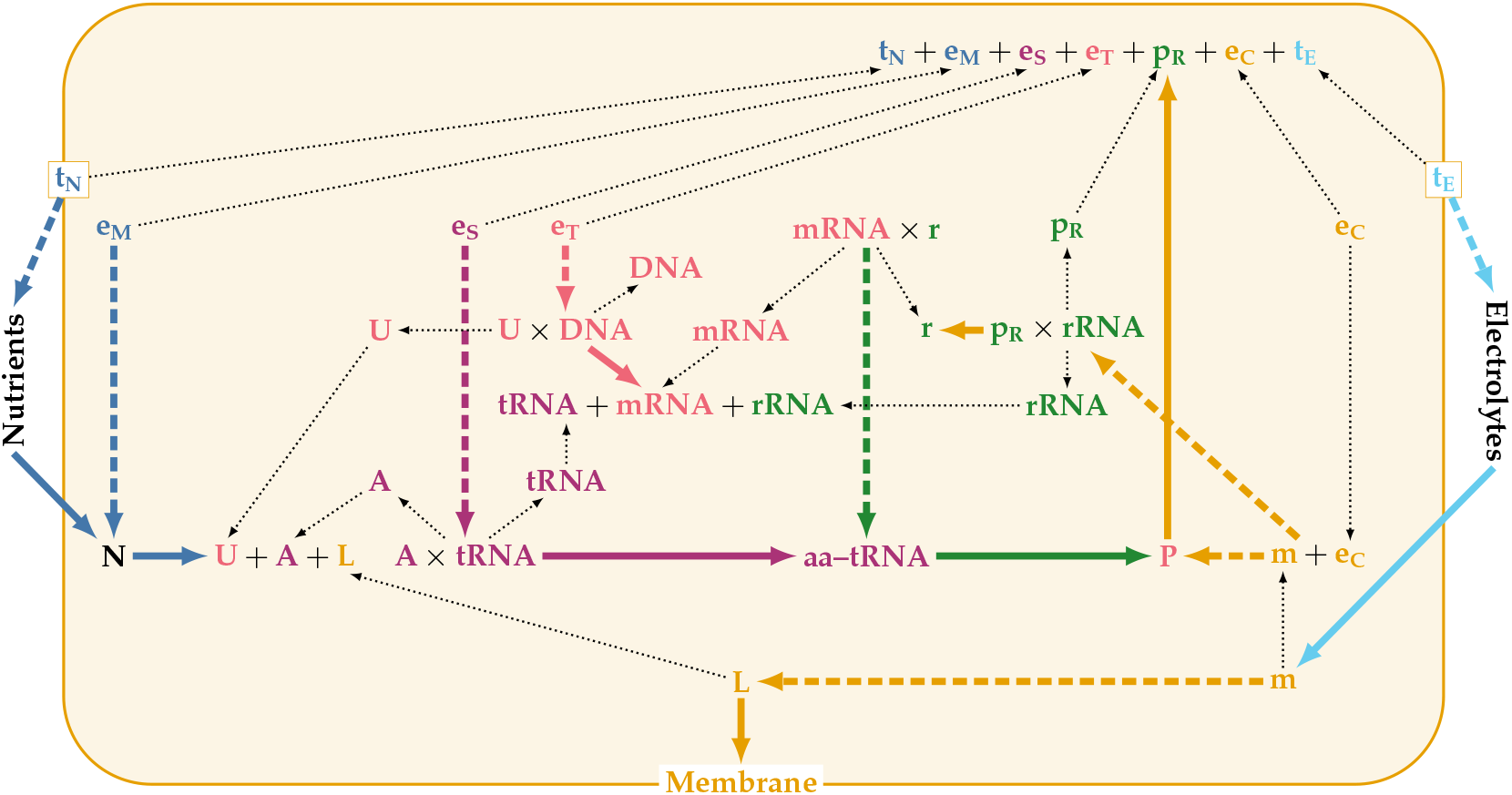
The (F,A) cell model: a graph-theoretic relational model of the self-manufacturing cell that is realised by the cell biochemistry in Fig. 10. *N*: nutrients, *U*: nucleotides, *A*: amino acids, *L*: lipids, DNA: deoxyribonucleic acid, RNA: ribonucleic acid, mRNA: messenger RNA, tRNA: transfer RNA, aa-tRNA: aminoacyl-tRNA, rRNA: ribosomal RNA, *P* = *P*_N_ + *P*_M_ + *P*_S_ + *P*_T_ + *P*_R_ + *P*_C_ + *P*_E_: non-functional, unfolded polypeptides, *r*: ribosomes, *t*_N_: nutrient transporters, *e*_M_: catabolic/anabolic enzymes of intermediary metabolism, *e*_S_: aminoacyl-tRNA synthetases, *e*_T_: transcription enzymes, *p*_R_: folded ribosomal proteins, *m*: intracellular milieu, *e*_C_: chaperones, *t*_E_: electrolyte transporters.

**Figure 12:**
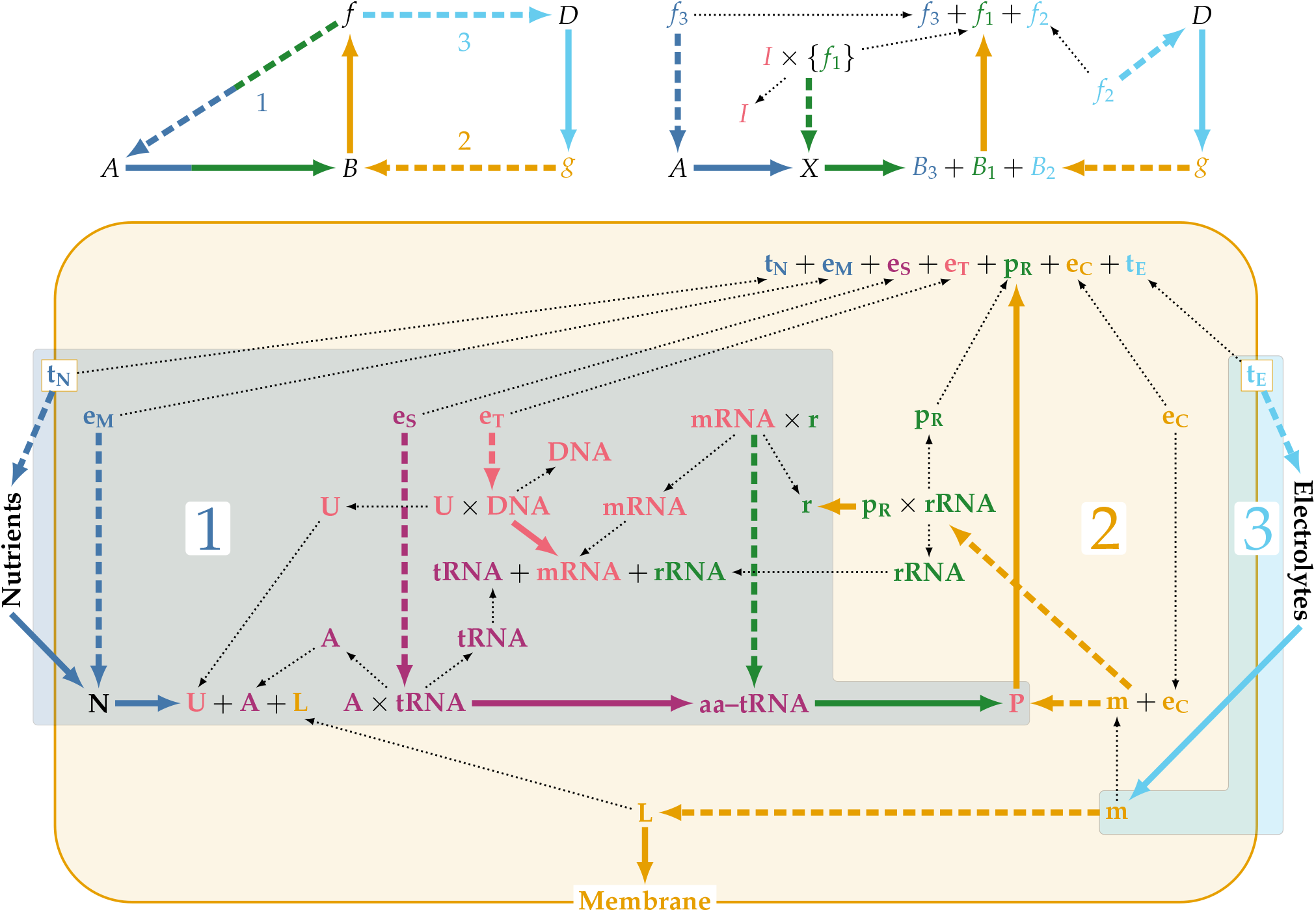
The closure of the three classes of efficient causes in the cell. (1) Metabolic enzymes *e*_M_, aminoacyl-tRNA synthetases *e*_S_, transcription enzymes *e*_T_ and ribosomes *r* catalyse the covalent metabolic reactions that transform the nutrients transported into the cell by *t*_N_ into the building blocks for the synthesis of polynucleotides (tRNA, mRNA, rRNA) and the set of unfolded polypeptides *P*, here equal to the direct sum *P*_N_ + *P*_M_ + *P*_S_ + *P*_T_ + *P*_R_ + *P*_C_ + *P*_E_. (2) A homeostatically-maintained intracellular milieu *m* and chaperones *e*_C_ drive the non-covalent, supramolecular folding and self-assembly of (i) polypeptides *P* into functional enzymes and transporters, and (ii) ribosomal rRNA and ribosomal proteins *p*_R_ into functional ribosomes. The intracellular milieu also drives self-assembly of lipids to form and maintain the cell membrane, thereby distinguishing and closing the cell from its surroundings. (3) Electrolyte transporters maintain the electrolyte composition of the intracellular milieu. The top left diagram is Fig. 4c, showing how its three mappings correspond to the three sets of efficient causes in the (*F,A*) cell model. The top right diagram is Fig. 9c, the starting point for the development of the (*F,A*) cell model.

The legends of Figs. 10, 11 and 12 are self-explanatory and provide all the necessary details. Table 1 provides a key to the use of colours in the (*F,A*) cell model diagrams.

**Table 1.** Key to the colour coding used in the (*F,A*) cell model diagrams and figure legends. (For interpretation of the references to colour the reader is referred to the web version of the article.)

| Colour | Process |
| --- | --- |
| Blue | Nutrient transport/intermediary metabolism |
| Green | Polypeptide synthesis |
| Red | Sequence information transfer |
| Violet | Genetic code instantiation in aminoacyl-tRNAs |
| Orange | Folding/self-assembly of polypeptides |
| Cyan | Electrolyte transport |

With respect to the model diagram in Fig. 11 a few comments are in order. First, as explained in Hofmeyr (2017), DNA not only serves as a template (formal cause) for mRNA, tRNA, rRNA, as well as for a host of non-coding RNAs involved in regulation, but also for its own enzyme-catalysed repair and replication. Furthermore, in eukaryotes the directly-transcribed pre-mRNA is processed into mature mRNA by adding a 5’ cap and a 3’ poly-A tail followed by the splicing out of introns and the possible rearrangement of the exons by means of alternative splicing. Adding all these processes to the model diagram would complicate it needlessly, but could be done by simply adding more enzymes and the spliceosome without it having any effect on the logic of the diagram. Transfer RNA, rRNA and the non-coding RNAs also need to fold into functional, threedimensional structures and, just as for polypeptides, the intracellular milieu is the driving force. Again, indicating this on the diagram would make it very complicated, so it is assumed. In fact, no matter how much the diagram is complicated by adding more features, the functional organisation remains unaltered as three sets of interlinked efficient causes: covalent catalysis by enzymes, supramolecular chemistry driven by the intracellular milieu, and maintenance of the intracellular milieu by membrane transport. The mapping *e*_M_: *N* → *U* + *A* + *L* represents the whole of intermediary metabolism, the hugely complicated enzyme-catalysed reaction network the diagrammatic representation of which adorns the walls of biochemistry lecture rooms worldwide.

Another factor, also discussed in Hofmeyr (2017) and impossible to depict on the diagram, is the contribution that metabolites, proteins and nucleic acids make to the homeostatic maintenance of the properties of the intracellular milieu, not as functional components but as chemical structures. The high protein concentration is the main contributor to molecular crowding, while inorganic and organic phosphates and proteins act as buffers that maintain the intracellular pH near to 7.2. Nevertheless, the homeostatic maintenance of the intracellular electrolyte composition by membrane transporters remains the major contribution to what makes the intracellular milieu chemically so different from the external environment (Hofmeyr, 2017) and can therefore be rightly regarded as its main efficient cause.

## 7. The manufacturing system in the cell

The relational diagram in Fig. 8d depicts an abstract fabrication-assembly manufacturing system. Fig. 13 shows its biochemical counterpart, which is embedded in the cell model in Fig. 11. The figure legend explains the role of the different components. In essence, the cell manufacturing system is a combination of a 1D fabricator and a 1D-to-3D assembler.

**Figure 13:**
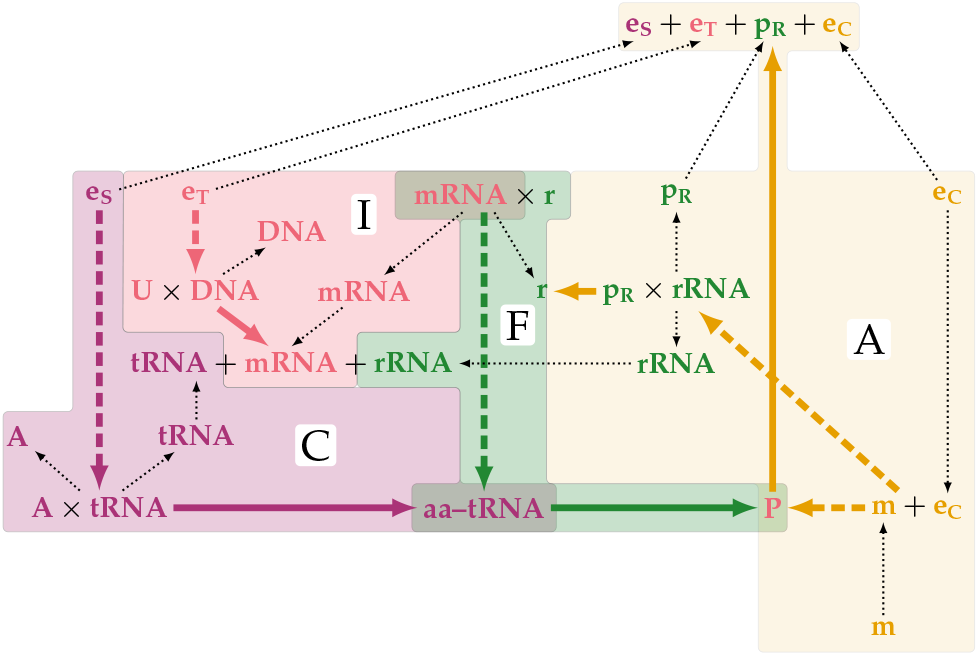
The manufacturing system in the cell. Block I represents the flow of sequence information (DNA to mRNA) through transcription catalysed by RNA polymerases (*e*_T_), block C the decoding system through the anticodon-carrying aminoacyl-tRNAs produced by aminoacyl-tRNA synthetases (*e*_S_), block F translation by the ribosome *r* (the fabricator) of the codon sequences in mRNAs into the amino acid sequences of the corresponding polypeptides *P*, while block A (the assembler) represents the intracellular milieu (*m* + *e*_C_) that drives both folding of polypeptides into functional enzymes and ribosome self-assembly from rRNA and folded ribosomal proteins *p*_R_.

Fig. 13 differs from the abstract manufacturing system in Fig. 8d in two respects. First, the decoding mechanism from codon in mRNA to amino acid in the polypeptide is made explicit by block c of the diagram, while it is implicit in the abstract manufacturing system. The rules of the genetic code, which is a mapping from three-letter codons to amino acids, are implemented by 20 aminoacyl-tRNA synthetases (*e*_S_) that couple anticodon-carrying tRNAs to their respective amino acids to form charged tRNAs (aa-tRNAs), the molecular adaptors that instantiate the genetic code. Second, while the fabricator *f*_1_ in Fig. 8d, given its own set of descriptions *I*_1_, can fabricate its full set of components *B*_1_, the ribosome can fabricate only its unfolded polypeptide components (*P_R_* ∈ *P*). The rRNA components of the ribosome have to be transcribed from DNA by RNA polymerases (*e*_T_) before ribosomes are assembled from folded ribosomal proteins *p*_R_ and rRNA. Because the ribosome cannot fabricate all of its components, let alone fold and assemble them, it cannot, as is often claimed, manufacture itself.

## 8. Realisations of the (*M,R*)-system in the cell

One vexing aspect of (*M,R*)-systems is the uncertainty about what counts as metabolism and what as repair. Fig. 14 captures possible realisations.

**Figure 14:**
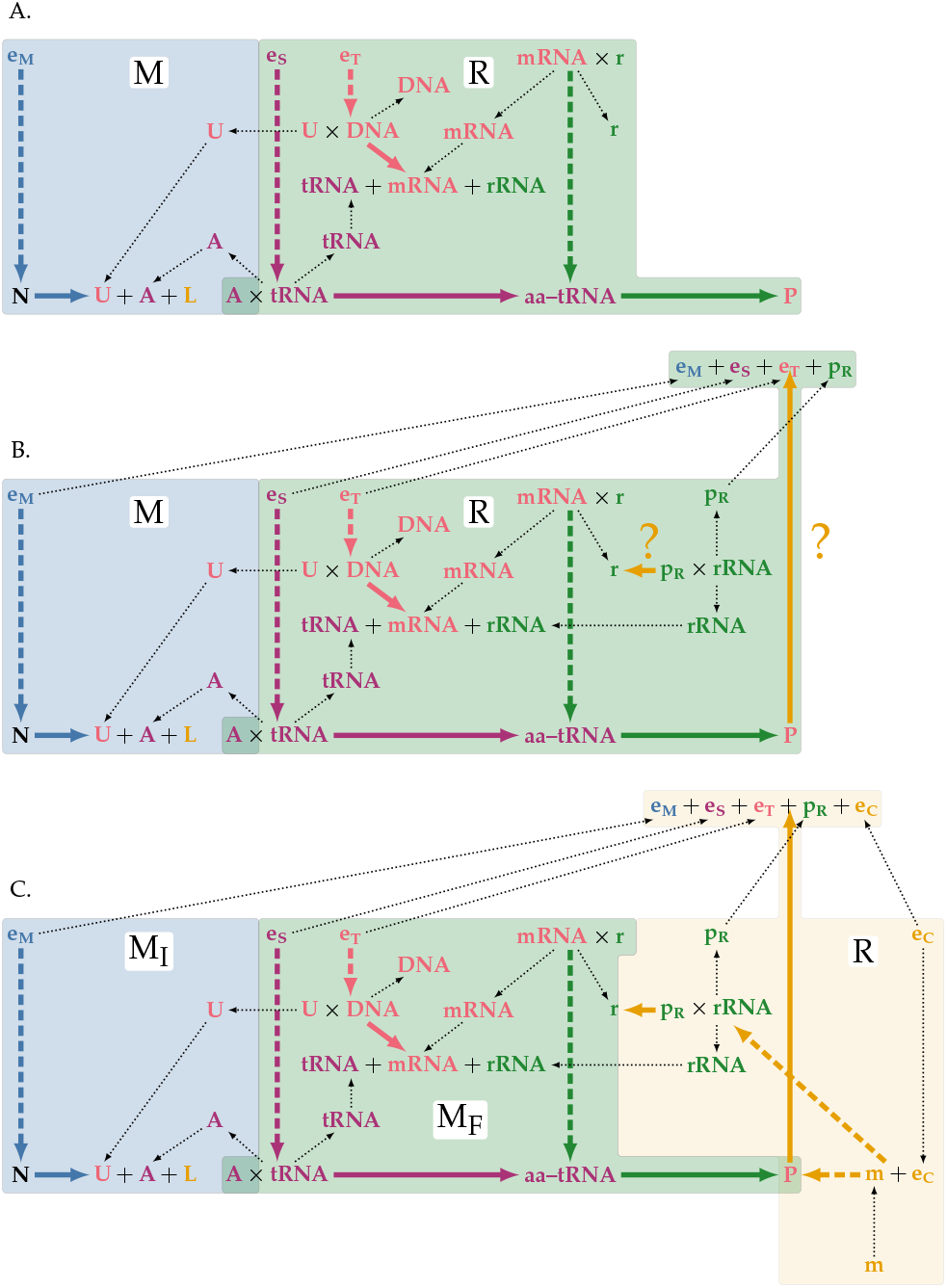
Possible realisations of metabolism (M) and repair (R). (A) Although the repair system *R* fabricates the primary structures *P* of the efficient causes of *M* (*e*_M_, *e*_S_, *e*_T_), they are not functionally entailed. (B) The repair system *R* produces functional enzymes, but the solid orange arrows that functionalise *P* and *r* have no efficient causes and are therefore still unentailed, hence the question marks. (C) The metabolic mapping *M* is a composition of intermediary metabolism *M*_I_ and biopolymer synthesis *M*_F_. The repair system *R* (the intracellular milieu) functionalises enzymes, transporters, ribosomal proteins, rRNA and ribosomes through supramolecular, non-covalent folding and self-assembly.

A first approximation is that intermediary metabolism provides the building blocks for the synthesis of biopolymers at the level of primary structure. This is captured by Fig. 14a, which is equivalent to Fig. 3b without *g* and the dashed arrow to *B*. In both Fig. 14a and 14b *B* represents amino acids as the intermediates between intermediary metabolism and polypeptide synthesis. The obvious problem in Fig. 14a is that the enzymes and ribosomes are not functionally entailed, which clearly discounts this interpretation. In an (*M,R*)-system the repair process must produce functional enzymes as in Fig. 14b, but here the question arises as to the efficient cause of the functional entailment of the enzymes and ribosomes. Fig. 14c shows the logical conclusion, namely that metabolism actually comprises the whole enzyme-catalysed network of covalent metabolic transformations (the composition of intermediary metabolism *M*_I_ and biopolymer synthesis *M*_F_) leading to the synthesis of the primary structure of biopolymers such as polypeptides and polynucleotides. Repair then comprises the functionalisation of these biopolymers through folding and self-assembly driven by the intracellular milieu. Louie (2017a,b) recently came to the same conclusion when he defined metabolism as material entailment and repair as functional entailment. In Fig. 14c material entailment (the solid blue and green arrows) is exactly equivalent to the covalent chemical transformations of metabolism, while functional entailment (the solid orange arrows) is the repair process that comprises folding and self-assembly into functional biopolymers. What Fig. 14c elucidates is that the efficient cause of functional entailment (dashed orange arrows) is the intracellular milieu and chaperones. This conception of metabolism and repair is therefore fully compatible with fabrication and assembly: the (*F,A*) cell model is in all respects an (*M,R*)-system. But, instead of Rosen’s (*MR*)-system with replication, we have an (*M,R*)-system with *functional context invariance* ensured by mapping *t*_E_ in Figs. 11 and 12.

**Figure 15:**
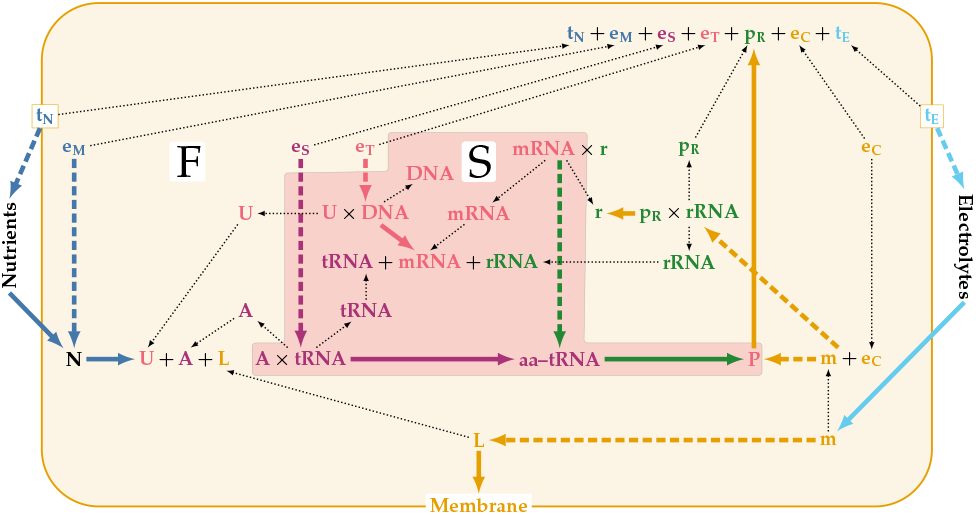
Pattee’s (1980) symbol-function (S,F) distinction bridged by the symbol-folding transformation as indicated by the solid orange arrows.

## 9. The epistemic cut between symbol and function

It was only after I developed the (*F,A*) cell model that I realised that it could be used to visualise other key concepts that have helped us understand what distinguishes life from non-life. For example, the metabolism-repair split in Fig. 14c coincides with Pattee’s (2001) primeval epistemic cut in the cell between symbol and function depicted in Fig. 15.

The epistemic cut describes a complementarity between a discrete symbolic (S) and a continuous dynamic functional mode (F), analogous to the digital-analog code duality of Hoffmeyer and Emmeche (1991). An epistemic cut is required to separate subject-object and symbol-function (or symbol-matter) distinctions. Pattee’s contention is that the “most primitive epistemic cut happened at the origin of life which separated the individual cell’s genetic *informational* constraints from the objective lawful dynamics it controls” (Pattee and Raczaszek-Leonardi, 2012, p. 5). This primitive epistemic cut is bridged by what Pattee (1980) called the *symbol-folding transformation*, the solid orange arrows in Fig. 15 (excluding the lipid to membrane arrow). Pattee is at pains to point out that the molecules that instantiate symbols comprise not only DNA and mRNA but also the unfolded polypeptides produced by ribosomal translation of mRNA; they are symbols in that they all only carry sequence information (Pattee, 1980).

## 10. The genotype-ribotype-phenotype ontology

Years before the advent of biosemiotics and organic codes, Barbieri (1981) introduced his ribotype theory of the origin of life. For ancestral systems Barbieri used the term ribotype to comprise all *ribosoids*, defined as molecules containing ribose. These were mainly RNA or complexes of RNA and peptides, and he defined the ribotype as the ‘collective of all ribosoids of an organic system’ (Barbieri, 2003). More precisely, Barbieri proposed that the ancestral systems evolved into triadic systems made of ribogenotype-ribotype-ribophenotype. Later on the ribogenotype evolved into genotype and the ribophenotype into phenotype, and it was this subsequent differentiation that gave origin to the first modern cells. Importantly, the ribotype theory confers on the ribotype a status that is ontologically differentiated from the genotype and phenotype.

In my attempt to visualise the modern ribotype with the (*F,A*) cell model the question arose as to the status of mRNA, which, although a form of RNA, should, as a carrier of sequence information, logically be classed together with DNA in the genotype. Since mRNA contains ribose, the original definition of the ribotype as all molecules containing ribose can therefore not apply to the modern cell. However, Barbieri suggests that it is possible that mRNAs evolved from ancestral anchoring-RNAs whose function was to hold in place the transfer-RNAs for a long enough time to allow the formation of a peptide bond, i.e., a manufacturing function not a sequence information function. Nevertheless, in Fig. 16 I do not include mRNA in the ribotype block. Furthermore, monomeric ribose-containing metabolites such as ribose sugars and their derivatives, nucleotides and their phosphorylated derivatives, as well as ribose-containing co-enzymes such as NAD, NADP, FAD and coenzyme A clearly play a phenotypic rather than the ribotypic role of mediator between genotype and phenotype, a role that is neatly captured by the diagram. In the light of this, Fig. 16 shows the ribotype comprising only tRNAs, rRNA, ribosomal proteins and the assembled ribosome (a nucleoprotein). A comparison of Fig. 13 with Fig. 16 reveals the close relationship between the ribotype and the cellular fabrication system (Blocks C and F in Fig. 13).

**Figure 16:**
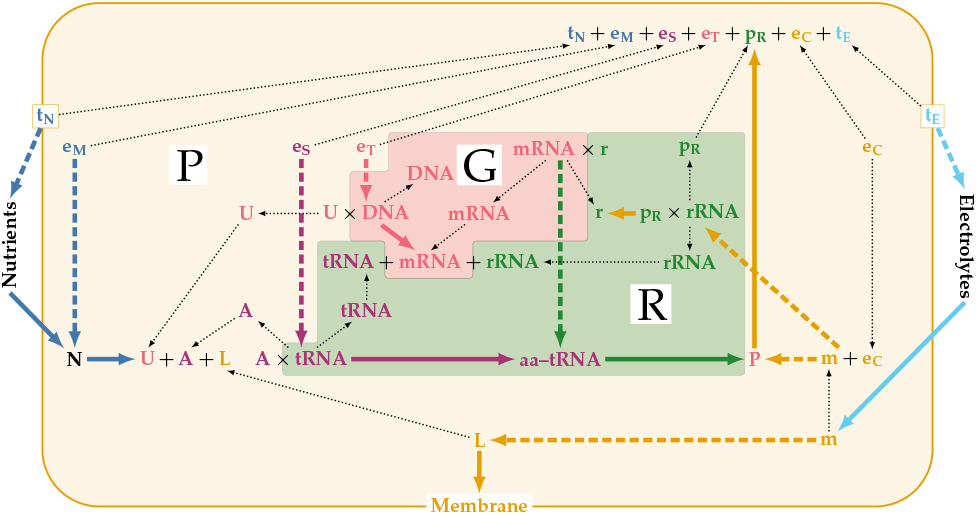
Barbieri’s (1981) genotype-ribotype-phenotype (GRP)-trinity.

## 11. Gánti’s chemoton

In 1971 Tibor Gánti (2003a,b,c) proposed a theory of life based on a model of a fluid automaton that he called the *chemoton*. The chemoton contains a metabolic cycle and an information cycle that interact to produce a self-assembling enclosing membrane in a third cyclic process. All three these cycles are described as autocatalytic in the sense of *A* + *X* → 2*A* + *Y*, a type of autocatalysis termed *reflexive autocatalysis* by Calvin (1969), although Reich and Sel’kov (1981) preferred the term *stoichiometric autocatalysis* to indicate its origin. Reich and Sel’kov (1981) emphasised that this type of autocatalysis is central to energy metabolism, in which an excess of energy equivalents (*A* in the reaction above) is by obtained by *sparking* a substrate *X* with a good leaving group.

Despite the many deep and important insights that came out of Gánti’s research, the chemoton has a serious deficiency: to isolate it from the network of spontaneous massaction reactions in which it is embedded its reactions must operate on a timescale orders of magnitude faster than the rates of the side reactions. This is kinetic isolation, not the topological isolation achieved by membrane enclosure, and can only be achieved by specific catalysts without which nothing prevents the chemoton’s intermediates dissipating into side reactions. The amplification achieved by autocatalysis does of course result in growth in the number of chemoton molecules in the system and a concomitant increase in reaction rates through mass-action, but this applies equally to the rates of the side reactions and therefore does not solve the problem. This problem of side reactions is a matter that I alerted one of the authors of Cornish-Bowden and Cárdenas (2020) to, and which they mention and acknowledge in their review. The reason for referring to it here is that pruning the (*F,A*) cell model into three blocks that correspond to the three cycles of the chemoton provides a stark visual picture (Fig. 17) of the lack of both catalysts and the active agency of the intracellular milieu, or, more broadly speaking, a lack of functional entailment.

**Figure 17:**
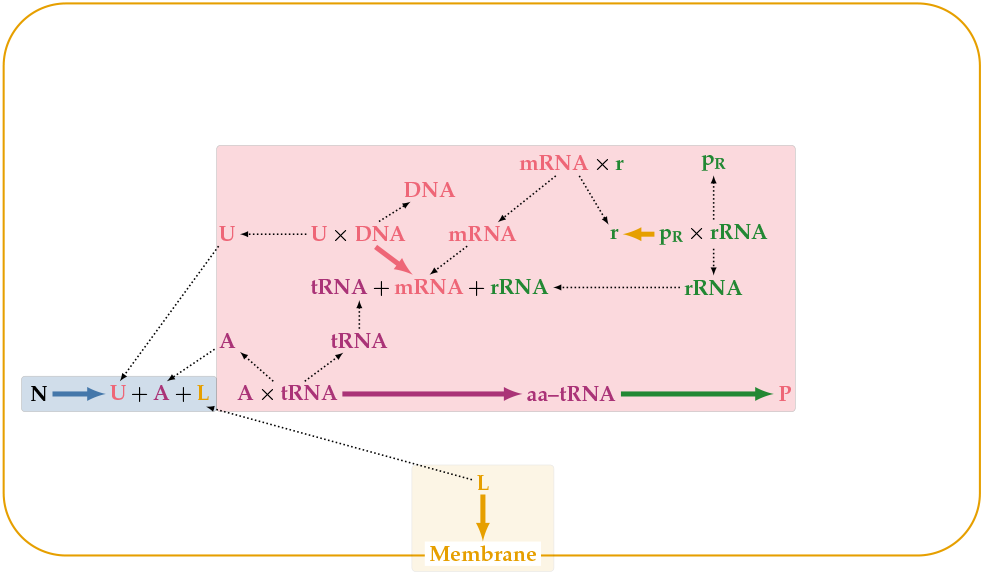
Gánti’s chemoton in the context of the cell. The metabolic, information and membrane components that correspond to the chemoton cycles are coloured blue, red, and orange respectively. The empty intracellular space indicates the absence of catalysts and the intracellular milieu.

## 12. Discussion

The problem of *realisation*, which stems from Rosen’s modelling relation, has long plagued his replicative (*M,R*)-system, both in his own hands and in those of many others. A modelling relation is a functorial relationship between a natural and a formal system (Rosen, 1991). Modelling is therefore the art of bringing the causal entailment structure of the natural system and the inferential entailment structure of the formal system into congruence with each other. In relational biological terms, using the category of sets and mappings, this means that objects in the natural system must map into sets in the formal system, and processes must map into mappings. If such a two-way relationship exists, the formal system is said to be a *model* of the natural system, and the natural system a *realisation* of the formal system. The usual approach in theoretical biology is to start with a natural system and seek to construct a model of it. Relational biology goes the opposite way: it starts with a model and seeks a realisation.

The latter is exactly what Rosen did with his replicative (*M,R*)-system. As recollected in Rosen (2000, Chap. 17) “I devised a class of relational cell models called (*M,R*)-systems (*M* for metabolism, *R* for repair). The idea behind these systems was to characterize the minimal organization a material system would have to manifest or realize to justify calling it a *cell*.” Furthermore, “It seemed to me (and still does) that one would not call a material structure a cell unless its activities could be partitioned into two classes, reflecting the morphological partition between nucleus (genome) and cytoplasm (phenome), and the corresponding functional partition between what goes on in cytoplasm (the *M* of the system) and what goes on in nucleus (the *R*).” From this he devised the (*M,R*)-system in Fig. 3b. Since mapping *g* was unentailed, he added a mapping *h* : *D* → *g* (Fig. 4b) and then further constrained the system by equating *D* with *B* (Fig. 4d). Now he had to find a way of pulling mapping *h* into the system in order to close it to efficient causation (although this way of phrasing it came much later). I particularly enjoy the understated way he describes his elation at figuring out how to do this: “Thus I was edified to discover that, under stringent but not prohibitively strong conditions, such replication essentially comes along for free, requiring nothing else but what is already in the diagram.” He then sketches out “the simplest way this can come about (although it is not the only way)” by introducing an invertible evaluation map to yield Fig. 4g, a mathematical sleight-of-hand that, while perfectly sound, has caused much confusion as well as misinterpretations that led to (all conclusively refuted) accusations of being wrong (see Cornish-Bowden and Cárdenas (2020)). What is, however, abundantly clear from his writings is that Rosen constructed the replication map purely on the basis of mathematical convenience, and not because he had some physical realisation in mind. Note that although his parenthetical remark acknowledges other possibilities of closing the (*M,R*)-system to efficient causation, he was clearly satisfied with his solution and so never explored other alternatives such as those in Fig. 4. *Since its inception the replicative (M,R)-system has remained the starting point of all elaborations of Rosen’s pioneering work*. In unkind moments I have come to think of it as akin to a black hole from whence, once sucked in, there is no escape.

The importance of finding biological realisations of (*M,R*)-systems is clearly stated in Rosen’s only attempt to address this issue (Rosen, 1971): “To make the theory of (*M,R*)-systems directly meaningful to a biologist, then, some point of contact must be found between that aspect of reality captured by the (*M,R*)-system formalism, and the kinds of system description employed in more conventional biological investigations.” After trying various approaches he admitted that he had not yet found a satisfactory solution to the problem.

The main problem throughout has been the realisation of (*M,R*)-systems in general, and more specifically the realisation of the replication mapping *h* in Fig. 4d. While Rosen’s idea of functional partitioning between metabolism and repair is perfectly sound (as shown in Fig. 14c), it has nothing to do with *M* being somehow associated with the cytoplasm, and *R* being associated with the nucleus. This idea was a red herring to start with since it excludes prokaryotes, which have no nucleus.

The choice of identifying *h* with *b* ∈ *B* brought about what is arguably the most serious realisation problem: on the one hand *B* is supposed to represent unspecified metabolic products from which enzymes *f* are synthesised, on the other at least some of its elements *b* must also be efficient causes that convert enzymes *f* to repair elements *g*. These dual roles for *B* acting as material cause of *f* and *b* as efficient cause for the production of *f* has proved difficult, if not impossible, to reconcile and has led to much, in my view mostly fruitless, discussion, e.g., Letelier et al. (2006); Cornish-Bowden and Cárdenas (2007); Cárdenas et al. (2010); Letelier et al. (2011); Mossio et al. (2009); Palmer et al. (2016); Zhang et al. (2016); Cornish-Bowden and Cárdenas (2021). With regard to the realisation of the replication mapping itself, renaming it to ‘organisational invariance’ (Letelier et al., 2006) or ‘coordination’ (Wolkenhauer and Hofmeyr, 2007) is of no help in identifying the cellular processes or agents that act as repairers of the repair system. Much has been made by the Santiago-Marseille group of their concept of ’systems with organisational invariance’, their translation of ‘(*M,R*)-systems with replication’ (Letelier et al., 2006). These are defined as systems in which *all* of the repair components of (*M,R*)-systems are regenerated, which is just another way saying that these systems are *clef*. In this meta-level sense I find the description ‘organisationally invariant systems’ a quite useful alternative to ’systems that are closed to efficient causation’, but then it must be consistently used only in that sense; ‘organisational invariance’ cannot at the same time serve as the name for one particular mapping in the system. Another suggested realisation of the replication mapping is ‘one gene–one enzyme’ (Louie, 2009). However, a gene is not an efficient cause: as sequence information, it is via transcription to mRNA the formal cause of an enzyme.

Nevertheless, an important finding, already hinted at by the dual roles that elements of *B* would have to play in the replicative (*M,R*)-system, came from Cornish-Bowden and Cárdenas’s (2007) analysis of their models of simple (*M,R*)-systems. They found that closure to efficient causation could only be achieved if some catalysts were assumed to fulfil more than one role, i.e., multifunctionality was required to ensure closure. For simple (*M,R*)-systems they suggested that the presence of ‘moonlighting proteins’ as defined by Jeffery (2003) could suffice; in this definition moonlighting proteins are single proteins with multiple functions that are not because of gene fusions, splice variants or multiple proteolytic fragments. However, they recognised that “the number of ‘moonlighting’ proteins known is rather small and falls far short of what is actually needed to create and (*M,R*)-system of significant size.” In the light of this they suggested protein synthesis as a possible source of multifunctionality in real organisms: using the sequence information in DNA, the small number of molecules that comprise the protein synthesis apparatus can produce all the proteins the cell needs. As they pointed out: “This can be regarded as multifunctionality on a very large scale”.

All of the above mentioned realisation problems are avoided by the (*F,A*) cell model in that all its mappings have clear realisations in terms of biological processes. The multifunctionality ‘on a very large scale’ is achieved by solving the problem of the implied infinite regress in the mapping *f*_1_ and its codomain *B*_1_ in Fig. 5a: fabricator *f*_1_ in Fig. 8 combines with a set of descriptions (formal causes) *I_i_* with *I*_i_ ∈ *I*, each individual description providing the information for the fabrication of a member of the set of components *B_i_*, which is then assembled into a mapping *f_i_* by assembler mapping *g*. In the (*F,A*) cell model the fabricator is the ribosome, which can synthesise as many polypeptides as the DNA codes for by using the transcribed mRNAs as freestanding formal causes. Cornish-Bowden and Cárdenas’s (2007) were therefore correct in their identification of protein synthesis as the probable source of multifunctionality in real organisms, although they could not relate that back to the replicative (*M,R*)-system, the relational structure of which does in fact not allow it.

In the (*F,A*) cell model the repair component is the intracellular milieu, and its active homeostatic maintenance by electrolyte transport (equivalent to its regeneration) is part and parcel of the model. In the (*F,A*)-system in Fig. 5 mapping *f*_2_ is the equivalent of Rosen’s replication mapping, while in the cell model in Figs. 11 and 12 it is mapping *t_E_*. Both these mappings ensure *functional context invariance*. In Hofmeyr (2007) I tried to capture the importance of this concept in the mantra “Nothing in an organism makes sense except in the light of functional context”, a systems biological counterpoint to Dobzhansky’s (1973) famous “Nothing in biology makes sense except in the light of evolution”.^8^

A bird’s-eye view of the causal entailments in the (*F,A*)-system and the (*F,A*) cell model shows that, while it is closed to efficient causation, it is open to both material causation and to the freestanding formal causes of the products of the fabricator. The openness to material cause ensures that the system is thermodynamically open and the openness to formal cause ensures that it is informationally open. Being thermodynamically open means that the system can act as a dissipative structure operating far from equilibrium (Nicolis and Prigogine, 1977), while being informationally open means that the functionality of the system (its efficient causes) can adapt and evolve through changes in the freestanding formal cause. The material causes also represent the so-called *admissible inputs* that have to be provided by the external environment of the cell (Giampietro et al., 2012).

Ever since I realised how important formal cause is for understanding cellular self-manufacture, it has puzzled me why other Rosen-inspired researchers have discounted formal cause in their analyses of (*M,R*)-systems, some, as I noted in a previous section, explicitly regarding it of minor importance for understanding metabolic closure. The only reason I can think of is that they misunderstand what formal cause actually is. Cárdenas et al. (2018) state that “the formal cause of an element is a definition of its role in a system: in a metabolic context, we call glucose 6-phosphate a metabolite because it is an intermediate in a metabolic pathway, in this case the harnessing of energy from glucose.” Similarly, Cornish-Bowden and Cárdenas (2020, 2021) state that the formal cause of glucose-6-phosphate (G6P) is that it is made as a glycolytic intermediate. In an earlier article they equate the formal cause of G6P to its status as a product of an enzyme-catalysed reaction (Cornish-Bowden et al., 2007). All of these statements assign G6P to an equivalence class in which it is indistinguishable from the other members in the sense that they are share the same formal cause. This is exactly the opposite of what formal cause should explain, namely what is it that makes G6P different from all the others in the equivalence class: it should tell us what it is to be G6P and not any other glycolytic metabolite or any other reaction product. Their interpretation amounts to saying that the formal cause of David is that it has been made as a statue in Michelangelo’s oeuvre of sculptures, or its status as a product of Michelangelo’s sculpting. If this were correct, then all Michelangelo’s statues would have the same formal cause, and, similarly, all glycolytic intermediates and all products of enzyme-catalysed reactions. In fact, all Michelangelo’s statues have the same *efficient* cause, namely Michelangelo. It is the combination of Michelangelo and his prior conception (formal cause) of each individual statue that distinguishes them from each other. Consider the formal cause of a particular polypeptide: in their definition its formal cause would be that it is made as a product of ribosomal protein synthesis, which does not distinguish it from other polypeptides. In truth, what it is *to be* that particular polypeptide inheres in its unique amino acid sequence, which is the actualisation of its formal cause, its mRNA.

I also find it difficult to understand why the seminal work of John Von Neumann (1951, 1966) on the logic of self-reproducing automata and that of Howard Pattee on the symbol-function distinction (the epistemic cut) and the bridging symbol-folding transformation (Pattee and Raczaszek-Leonardi, 2012, p. 158) has found no traction in all of the discussions and analyses of (*M,R*)-systems. The extensive literature on biological autonomy at least takes Pattee’s work seriously (Moreno and Mossio, 2015). The (*F,A*)-system makes it clear that these concepts are indispensable for our understanding of life and must therefore form part of any theory of life. In Cornish-Bowden and Cárdenas’s (2020) review, Von Neumann’s work does not even warrant a proper discussion and is dismissed on the grounds that “self-fabricating machines have yet to be built, and for the moment the gap between machines and organisms appears to be unbridgeable”. Is this a valid reason for not acknowledging his ground-breaking logic of self-reproduction? Similarly, the whole of Pattee’s work is discounted; he is only mentioned as describing “the evolution of Rosen’s thought over the years until Life Itself (Rosen, 1991) appeared”.

To what extent does the (*F,A*) cell model address the four core challenges that all organisms have to overcome if they are to persist as autonomous entities (Table 2)? Clearly it explicitly addresses the solution to the first of these challenges, namely how the problem of being built from fragile components with short lifetimes compared to the lifetime of a cell is overcome by autonomous self-manufacture. The model acknowledges the second challenge by incorporating the efficient cause of the self-assembly of the cell membrane from intracellularly produced lipids. According to Maturana and Varela (1980), together these two properties define the cell as being *autopoietic*.

**Table 2.** Four core challenges to the persistence of the organism.

| Challenge | Solution |
| --- | --- |
| Built from fragile components | Functional organisation that allows autonomous self-manufacture |
| Distinguish itself from the rest of the world (individuation) | Self-bounding with a semi-permeable membrane |
| Cope with environmental fluctuations during its lifetime | Sensing the environment and adaptive restructuring of cellular functionality within genomic constraints |
| Cope with environmental changes over generational time-scales | Evolutionary adaptation through descent with modification (natural selection) |

The functional organisation of self-manufacture allows the cell to respond to environmental fluctuations, the third challenge, by making it possible to reconfigure its catalytic repertoire within the functional space allowed by genomic constraints. What is missing from the (*F,A*) cell model are the mechanisms that allow the cell to sense such changes in its external environment; as depicted in Fig. 11 the cell model is ‘blind’ to its *Umwelt* (Uexküll, 2001). However, adding the synthesis of the membrane receptors and components of their signal transduction networks, all of them proteins, is easily accommodated by the manufacturing system since their formal causes are of course encoded in the DNA, just as those of all other functional components of the cell. The translation by the sensing system of external molecular signals into intracellular effects depends on the signal transduction code, without which the cell cannot be said to be alive (Barbieri, 2015).

The last challenge is not a challenge to the persistence of an organism, but rather to the persistence of its lineage over generational timescales in the face of environmental change. Here an organism needs to be able to self-reproduce, which means being able to grow, replicate its DNA and divide, all processes that also require the ability to self-manufacture. Being informationally open, the cell model allows for changes in the DNA, its primary formal cause, which bestows on any lineage of organisms the ability to adapt to environmental changes over generational timescales through natural selection.

A distinguishing feature of the (*F,A*)-system is the absolute need for an actively-maintained, stable internal context that ensures that the components of the system can fulfil their individual functions efficiently, a clear example of a whole-system constraint that acts through downward causation (Auletta et al., 2008; Farnsworth, 2017; Farnsworth et al., 2017; Ellis, 2020). In the living cell this translates as the functionalisation of polypeptides by the homeostatically-maintained intracellular milieu. If the properties of the intracellular milieu should change to the degree that folding and self-assembly become impaired, the other efficient causes in the cell will lose their function and their replacement becomes impossible, which, in turn, makes it impossible for them to repair the intracellular milieu. Over time the functionality of the whole system will collapse. This collectively impredicative relationship between the efficient causes of the folding transformation (the intracellular milieu) and the efficient cause of homeostatic maintenance of the intracellular milieu (electrolyte transporters and chaperones, both functionalised by folding) is captured by the hierarchical cycle in Figs. 4c, 5a and 9.

The hierarchical causal cycle of the functionality of the creators and maintainers of internal context being determined by that very context exists at many levels of the organisation of the living. Here I have considered what is arguably the lowest of these levels, but the requirement for an enabling context created and maintained from within the system scales from the cell right up to human societies. I leave it to the reader to consider how such a hierarchical cycle operates at the level of, for example, a multicellular organism, an ecosystem, an economy, a human organisation or a society. Such reflection is important, because we are confronted daily with the consequences of the breakdown of hierarchical cycles: cancer destroying bodily homeostasis, climate change and human intervention destroying ecologically constructed niches, a toxic organisational culture making it impossible for the employees of a firm to do their job, societies that collapse because their cultural norms are not maintained, liberation movements unable to govern because of their inability to shift their internal organisational culture to one commensurate with serving their people and not themselves. What I have demonstrated in this article is that the living cell has learnt to avoid this trap: it has figured out Life’s Trick, namely how to create and maintain a stable functionalising context that enables the harnessing of the supramolecular chemistry of the symbol-folding transformation. For us the cell’s lesson is to realise that our functionalising contexts have agency, and should therefore be continuously monitored, cherished and actively maintained from within by the members of our organisations and societies.

## Acknowledgements

I thank Aloisius Louie, Johannes (Yogi) Jaeger, Marcello Barbieri and Andrei Igamberdiev for fruitful discussions on the topic of this article and related issues. As with previous work, Aloisius graciously clarified some mathematical aspects of relational biology. Through his close reading of and insightful commentary on the article, Keith Farnsworth made a huge contribution to the clarification of a number of aspects. Part of this work was initiated while on a Fellowship of the Wissenschaftkolleg zu Berlin in 2014/15. The work as a whole is the culmination of a project made possible by the 2002 Harry Oppenheimer Fellowship Award, which allowed me to embark upon a new research path. In the end my greatest thanks must go to the late Robert Rosen for his incredible pioneering work on the question of what makes living things different from non-living ones. I like to think that he would have appreciated this article.

## Footnotes

1 A mapping that *directly* entails itself is an impossibility in Cantorian set theory, and therefore impossible in the category **Set** and any concrete category in general (Louie, 2009, 2013).

2 For the present purpose equating final cause with function will suffice, but this is philosophically contentious. For different views of the role of final cause, function and purpose in biology see, for example, Cummins (1975); Mossio and Bich (2014); Farnsworth (2017); Farnsworth et al. (2017); Cooper (2020).

3 Although Rosen (1991) is most often cited here, Rosen (1972) provides his most detailed mathematical description of this system; see also Letelier et al. (2006) and Louie (2009).

4 Here the simplifying assumption is that the reaction rate in the absence of enzyme is negligible, so that the reaction product for all intents and purposes does not form without enzyme present. A more subtle approach would recognise the efficient cause of any enzyme-catalysed chemical reaction such as A → B asa combination of catalytic action, which speeds up the reaction rate, and the thermodynamic driving force quantified by the disequilibrium ratio *ρ* = Γ/*K*_eq_, where Γ is the mass-action ratio of concentrations [B]/[A] and *K*_eq_ the equilibrium constant of the reaction. Similarly, the formal cause is a combination of, on the one hand, the specificity of the enzyme, which ensures that the active site of the enzyme ‘chooses’ to bind and convert to B only substrate A and not any of the other compounds in the mixture, and, on the other, the intrinsic propensity of A to transform chemically into B.

5 Hofmeyr (2018) discussed the other possibility where formal and material cause associate with each other as separate entities in the mapping *f* :Σ × *A* → *B* in which *b* = *f* (*σ, a*) with *a* ∈ *A, b* ∈ *B* and *σ* ∈ Σ.

6 Hoffmeyer (2000, p. 177) agrees that by creating topological closure the cell membrane establishes “an inside-outside asymmetry, which is an absolutely decisive step because it opens the door to the semiotic world of communication and function; and thereby to the formation of an *individualized context space* or *agency*”.

7 https://reprap.org/wiki/RepRap, accessed 26/04/2021

8 Originally, in Hofmeyr (2007), I phrased it as “Nothing in an organism makes sense except in the light of context”, but I have since added the “functional” to make it clear that the context is active. In the light of the (*F,A*) cell model I would now even elaborate it as “homeostatically-maintained functional context”.

## Notes

### Competing Interest Statement

The authors have declared no competing interest.

